# Historical and Contemporary Management of Genetic Diversity in French Local Horse Breeds: Insights from Genomic Data

**DOI:** 10.64898/2026.10.21.760631

**Authors:** Y. Shakya, C. Bonnin, S. Danvy, B. Dumont - Saint Priest, E. Rousseaux, M. Sabbagh, E. Verrier, B. Servin, G. Restoux

## Abstract

The phenotypic and genetic diversity of horses worldwide today likely resulted from two main events: (i) domestication and, afterwards, (ii) breed creation. While many studies discuss the origin and domestication of horses, very few focus on the second important event, which has largely shaped the genetic architecture of horses at present. This study exhaustively examined all 18 local horse breeds from France, all of which are endangered by small census sizes, making them prone to suffer from genetic drift, loss of diversity, and inbreeding depression. While pedigrees are available, they are often incomplete or error-prone. Thus, we used molecular markers to reconstruct the history of French local breeds and to monitor the evolution of genetic diversity and its link to management practices. The study was based on 976 animals (≈50 per breed), spanning two time periods corresponding to the change in management from public stud farms to privatization. The samples were genotyped using a 10K SNP array dedicated to genetic resources characterization. French local horses can be classified not only phenotypically but also genetically into two categories: the light “territory” breeds and heavier “draft” breeds, with regular gene flow observed between the two groups that are linked to historical records. The evolution of kinship and inbreeding over the two time periods exhibited distinct patterns between the cohorts and across the breeds, reflecting various consequences of different management practices. To conclude, this study was the first to exhaustively depict the evolution of French local horses, while it also demonstrated the usefulness of genomic data in monitoring genetic diversity and evaluating the effectiveness of management programs.

## 1. Introduction

Horses today show great phenotypic and genetic diversity worldwide, which likely results from two main events: (i) domestication and, afterwards, (ii) breeds’ creation. The domestication of modern horses has long been debated, however, with modern sequencing tools and ancient DNA analyses, we now have strong evidence that modern horses descended from the DOM2 lineage (which diverged from the Botai horse around 35-55 thousand years ago) and were first domesticated in the Western Eurasian steppes, specifically the lower Volga-Don or Pontic-Caspian region, around the late fourth and early third millennia BC (Guimaraes et al., 2020; Orlando, 2020a; 2020b; Librado et al., 2021). As DOM2 horses spread across Eurasia, they replaced local lineages by introgressing with their maternal lines, as supported by the rich mitochondrial diversity in horses, not seen in other domestic species (Vilà et al., 2001; Olsen et al., 2006; Orlando, 2020b; Guimaraes et al., 2020; Librado et al., 2021). The domestication and companionship of horses enabled humans to conquer the world, leading to the interconnected world we live in today (Khadka, 2010). This occurred alongside the selection of horses for traits such as speed, stamina, appearance, and size, depending on the purpose they served in different parts of the world, along with changes in management, such as close breeding and the formation of isolated populations now referred to as breeds (Librado et al., 2021; Orlando, 2020b). Breed creation, especially over the last few centuries, has largely influenced the genetic architecture that was relatively stable for the 4000 years after domestication (Fages et al., 2019; Orlando, 2020a). While many studies discuss the origin and domestication of horses, very few focus on the second important event: breed creation, which reshaped today’s horse populations.

Despite being a major workforce in transforming the world through socio-economic development, the rise of mechanization has recently limited the use of horses to sports and leisure activities (Khadka, 2010; Cressent & Jez, 2013; Gosselin, 2023a). This has led to increased interest in a few international transboundary breeds, like Arabians and Thoroughbreds, and a rapid decline in the local horse populations by the 20th century (Khadka, 2010; OIE, 2022). A report from the United States showed that there were 27.5 million horses in 1910, which plummeted by 85% to 4 million by 1960 (‘USDA Horse Total, 1850-2012’). Similar cases of drastic population decrease were observed in Asia and Europe, leading to many local breeds becoming endangered (Verrier et al., 2015; Lepetz et al., 2021; OIE, 2022; Tim H, 2025). According to FAO’s 2024 data, there are around 57.6 million horses, representing 714 local breeds globally. Of these local horses, 302 are at risk: 89 are classified as endangered and 148 as critical, while the risk status for 346 breeds has not been reported (FAO, 2024). Lack of reports, mostly from developing countries, is largely due to the lack of characterization of local breeds, leading to an even worse picture of horse breed conservation (Khadka, 2010; *FAO*, 2024).

Although local horses are no longer used for traditional purposes in many parts of the world, they remain important animal genetic resources; they thrive in challenging climatic and geographic conditions, with remarkable resilience and adaptability to their local regions. These animals are invaluable assets and prerequisites for developing diversified agriculture and sustaining farming systems, land use, and maintenance, which could play an important role in the agroecological transition (Rzekęć et al., 2020). Their role in promoting cultural heritage and traditions, equestrian tourism, and biodiversity conservation is significant (Gosselin, 2023a). Additionally, novel utilities such as equine-assisted therapy offer promising benefits to mankind (Jez et al., 2013). Each of these local breeds is an important unit of conservation, and together they contribute to the overall diversity of a species once seen in ancestral horses. The loss of these breeds threatens not only their population but also the species and overall local biodiversity. Therefore, it is crucial to protect these local breeds, even though they may not currently hold high economic value; they represent a secure resource for future needs.

Conserving local breeds presents several challenges, including limited characterization, insufficient pedigree and phenotyping records, and small, isolated populations prone to inbreeding and genetic drift (FAO;SoW-AnGR, 2019; ERFP, 2021). As a result, preserving and maintaining them requires diligent, methodical steps and decision-making. The three major steps of conservation for local breeds are: discovery, securing, and sustaining (Sponenberg et al., 2019). With advances in genomic technology and molecular tools, molecular markers are emerging as rapid and cost-effective alternatives to pedigree information for characterization and management of the breeds (FAO, 2007). In recent years, single-nucleotide polymorphisms (SNPs) have been used widely, offering several benefits, such as cost-effectiveness, automated, straightforward allele scoring, and wider genome coverage, which provides greater accuracy in describing individual and breed relationships (FAO, 2007; Husien et al., 2024).

This study examines a case from France, a country recognized for its rich equine diversity and the historical significance of horses in its development, as evidenced in the arts and historical records (Lepetz et al., 2021). French local horses are categorized into two groups: heavier draft horses and lighter territory horses, with nine unique breeds in each category (SFET, 2022). Originating in specific regions and named after them, these breeds not only exhibit distinctive traits but also contribute to regional identities (Jez et al., 2013). These breeds have specific breed associations and organizations, and their management used to be ensured by national public bodies, but in 2015 a major restructuring was implemented that transitioned horse management, including breeding and stallion activities, to local and private authorities (*Avenir Des Étalons Nationaux*, 2014).

Since the early 20^th^ century, France has seen its horse population decline from 3 million to half a million by the end of the century, mostly affecting local or draft horses used for practical purposes (Leroy et al., 2009; Rzekęć et al., 2020). Today, all these breeds are considered at risk or endangered, some with alarmingly low numbers (Allier et al., 2023; Verrier et al., 2015). For instance, SFET reported 5 births in 2022 in Cheval-Corse, 25 in Barraquand, 31 in Castillonnais, etc. Several *in-situ* conservation and dedicated breeding programs have been implemented for equines in France (Leroy et al., 2009; Jez et al., 2013). However, these efforts are mostly based on pedigree information (Dubois & Ricard, 2007). Despite pedigree data being useful for preserving and managing some wild and livestock species, it can suffer from accuracy issues due to depth, missing information, and human error (Pirault et al., 2013; Galla et al., 2022). Additionally, many equine farms focus on producing sports horses, with local breeds typically raised by hobby breeders who manage small herds (Cressent & Jez, 2013). Consequently, information on the pedigree of these local horses is often limited, highlighting the challenges of genetic management (Pirault et al., 2013). Given these issues and recent changes in horse management, applying genomic tools could offer valuable insights into the evolution of genetic diversity and management of these breeds.

Our study aims to exhaustively document the origin of these French local horse breeds by tracing their histories and assessing their genetic evolution in response to recent management changes, considering both historical and recent perspectives to inform conservation decisions. The objectives of the study are to (i) reconstruct the evolution and relationships between breeds by assessing the neutral, spatial, and adaptive structure of French local horses, (ii) evaluate the level of within-breed genetic diversity, and (iii) assess the effectiveness of breed genetic diversity management between the two time periods marked by the change in management from nation-run stud farms to privatization.

## 2. Materials and Methods

### 2.1. Sampling and Genotyping

The blood samples were collected from the two major breed groups in France, namely draft and territory horses, each comprising nine breeds. Pedigree information was used to sample non-related individuals (i.e., full and half-sibs) and to be as representative of each breed as possible by sampling various families. The total sample includes 992 individuals born between 1982 and 2022, grouped into two time periods referred to as cohorts in the paper. Animals are grouped according to changes in the management system; hence, those born before 2015 are grouped as cohort 1 (393 samples), and those born after are grouped as cohort 2 (609 samples). The mean birth years for cohorts 1 and 2 are ∼2003 and ∼2016, respectively, indicating an approximate gap of 1.5 generations. The distribution of samples throughout the birth years is shown in Supplementary Figure S1. Following DNA extraction, genotyping was performed using the Affymetrix® Axiom™ IMAGE 001v2 Multispecies SNP array (Crooijmans et al., 2025) with a final number of 7912 reliable SNPs. The location of sample collection, along with the breed names, is presented in Figure 1. Details of these breeds, including names, sample sizes (total and across the cohorts), and locations in coordinates, are presented in Supplementary Table S1.

**Figure 1:**
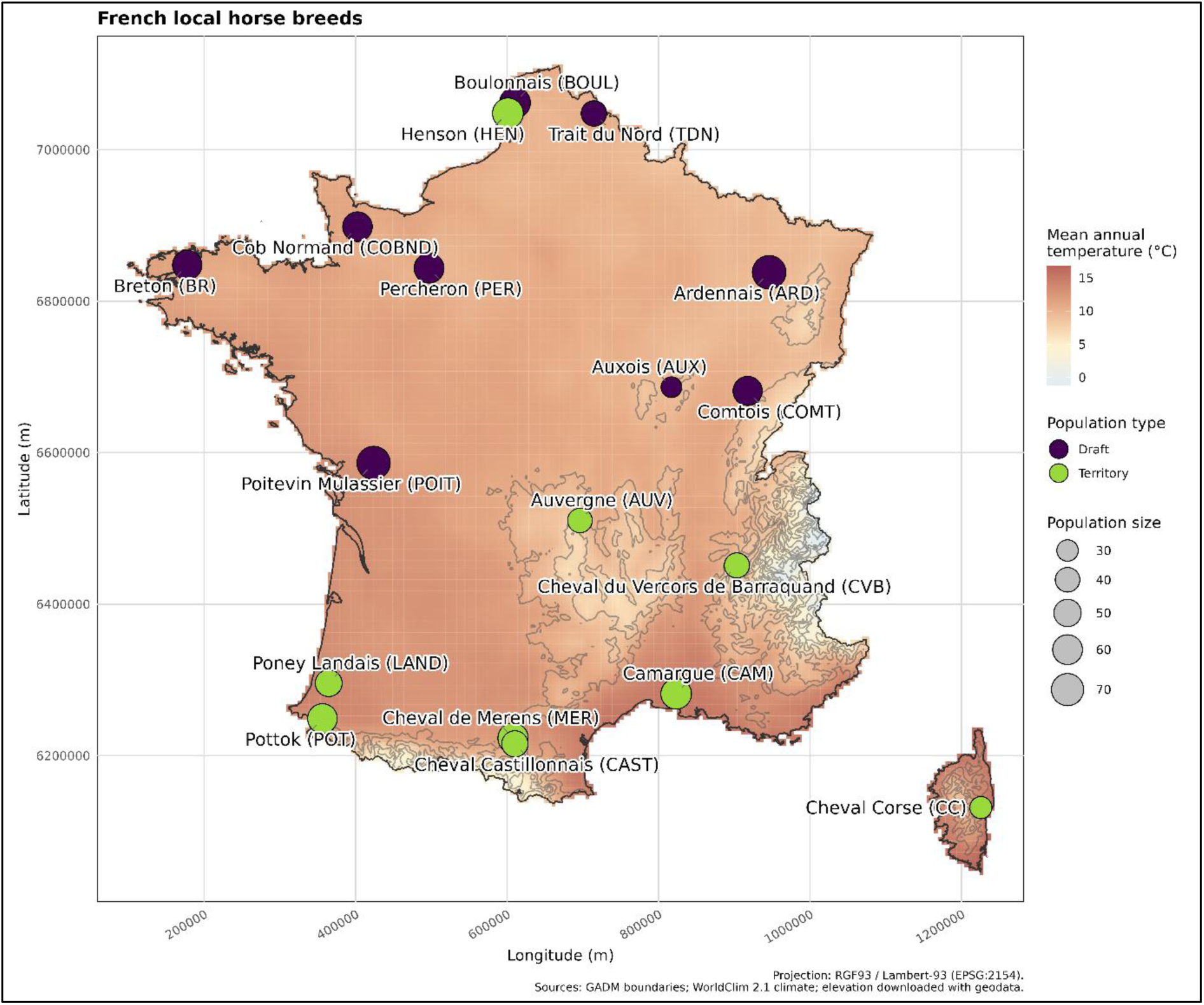
The circular points on the map mark the locations of French local horse breeds. The colour of the points indicates the breed group each breed belongs to, with purple for draft and green for territory, and the size of the point scales with the sample size. For the exact sample size refer to the Supplementary Table S1.

### 2.2. Quality Control and Datasets

General filtering applicable to all downstream analyses was conducted by removing SNPs located on chromosomes X, Y, and unidentified chromosomes, and those with call rates below 99%, resulting in a total of 7,669 SNPs. Twenty-six samples with a missing proportion (--mind) greater than 5% were discarded, resulting in a final sample size of 976 individuals (380 and 596 in cohorts 1 and 2, respectively). Based on the purposes of the analyses, two datasets were created: one without accounting for the cohorts, and the second with the breed names extended to “_I” for cohort 1 and “_II” for cohort 2. Furthermore, these datasets were filtered at both the global and breed levels to meet the needs of the different analyses with PLINK, version 1.9 (Chang et al., 2015; Purcell & Chang, 2019) (Supplementary Table S2).

### 2.3. Population Structure

Different analyses were used to provide a comprehensive view of the population structure of these local horses. First, a Principal Component Analysis (PCA) was performed as a preliminary step to summarise the data and examine population structure using PLINK with the *--pca* command. An unsupervised admixture analysis was performed with ADMIXTURE v.1.3 (Alexander & Lange, 2011) on a complete and balanced dataset to estimate the ancestry and obtain admixture fractions. The balanced dataset was obtained by subsampling 40 individuals from each breed, except Auxois (AUX) (28 individuals) and Cheval Corse (CC) (32 individuals). Subsampling was performed by removing individuals with the highest average kinship coefficients from the kinship matrix, recursively until the desired number of animals was retained. *POPKIN* R package (Ochoa & Storey, 2016, 2021) was used to compute the kinship coefficient with the function *popkin()*. K values ranging from 2 to 18 (the total number of breeds) were used to obtain cross-validation errors. The admixture graph was plotted using the R package AncestryPainter.V2 (Chen et al., 2024).

An identity-by-state matrix at the individual level was computed using the PLINK function ‘-- distance square ibs’. The dissimilarity matrix was converted to a neighbour-joining distance matrix using the *ape* R package (Paradis & Schliep, 2019) and an unrooted tree was plotted using the *ggtree* R package (Yu et al., 2017, 2018).

Pairwise-F_ST b_etween breeds was computed using the R package *DARTR* (Gruber et al., 2018), which utilises the method proposed by Wright (1949) and updated by Weir & Cockerham (1984) and an unrooted neighbour-joining tree was created using the R packages *ape* and *ggtree*. Moreover, to obtain a rooted tree, hapFLK (Fariello et al., 2013) was used to compute the Reynolds Distance (Reynolds et al., 1983).

Further, to obtain an in-depth understanding of the history of these populations in terms of population splits and gene flow, we used the software Treemix v1.1 (Milanesi et al., 2017; Pickrell & Pritchard, 2012; Zecca et al., 2019) which returned a maximum-likelihood tree (i.e., assuming genetic drift only) and added migration events.

### 2.4. Landscape Genomics

For each population, the breed association’s address was considered the “centre of origin” and used as a reference to obtain a single geographic coordinate in degrees from the French public web portal (https://www.geoportail.gouv.fr/), as shown in Supplementary Table S1. The coordinates were converted into a distance matrix using the R package geosphere (Hijmans, 2024) to test for spatial autocorrelation and isolation by distance (IBD). A Mantel test was performed between transformed genetic distances (F_ST/_1-F_ST)_ and the logarithm of geographic distances using the vegan R package (Oksanen et al., 2001; Diniz-Filho et al., 2013) separately for draft and territory breeds (Rousset, 1997). Henson from the territory group was removed from the analyses as it is a newly formed breed, which violates the IBD assumptions (Ishida, 2009; Malécot, 1948; Wright, 1949).

The climatic variables, elevation, land coverage data, and global soil data were retrieved from the WorldClim database v2.1 (Fick & Hijmans, 2017), the Shuttle Radar Topography Mission (SRTM) v4 (Jarvis et al., 2008), the ESA WorldCover data v100 (Zanaga et al., 2020), and the SoilGRIDS database v1 (Hengl et al., 2017) based on the coordinate information. Climatic variables include 19 standard bioclimatic variables at 10-minute (19 km) spatial resolution, averaged over 30 years (1970 – 2000); elevation at 3-second (90-meter) spatial resolution from the year 2000; land cover at 0.3-second (10-meter) spatial resolution for the year 2020; and global soil data at 30-second (∼1km) spatial resolution with the latest update in 2016. Throughout the article, we will refer to these variables together as environmental variables. To prevent collinearity among the environmental variables, we conducted backward variable selection, sequentially removing the variables having a variance inflation factor (VIF) greater than 10. The complete list of selected environmental variables is shown in Supplementary Table S3.

Redundancy and partial redundancy analyses were performed using the Vegan R package (Oksanen et al., 2001). For partial redundancy analysis, we used population structure and environments as covariates in the models, as shown in Table 1. For population structure, we used the admixture Q-matrix computed for a K=2 with the Tess3r package (Höhna et al., 2015), which accounts for both geographic and genomic structure. As we have single-location coordinates, hence a single value per environmental variable for the individuals within a breed, we did not opt for higher *K* values because fine-scale clustering accounting for overall structure greatly reduces the statistical power of detecting any variations explained by the environmental variables (Richardson et al., 2016).

Variance partitioning was performed to obtain the inertia explained by environment, structure, and the confounding factor between them. This latter term was obtained by subtracting the adjusted R-squared of the full model to the partial models of environment conditioned by structure and structure conditioned by environment, respectively (Capblancq & Forester, 2021). Lastly, since RDA does not permit missing genotype data, genotypes were imputed using the *snmf* function in the LEA package (Frichot & François, 2015). The formulas for the models are in Table 1:

**Table 1:** Full and partial RDA models and their *formulas*.

| Model | Formula |
| --- | --- |
| Full | $\text{Genotype}_{(n \times m)} \sim \text{Environment}_{(n \times p)} + \text{Structure}_{(n \times q)}$ |
| Environment | $\text{Genotype}_{(n \times m)} \sim \text{Environment}_{(n \times p)} + \text{Cond}(\text{Structure}_{(n \times q)})$ |
| Structure | $\text{Genotype}_{(n \times m)} \sim \text{Structure}_{(n \times q)} + \text{Cond}(\text{Environment}_{(n \times p)})$ |

With, $n$ : number of individuals $m$ : number of loci $p$ : number of environmental variables $q$ : number of $K$ -ancestry

### 2.5. Genetic Diversity

#### Heterozygosity, F-statistics, mean MAF, and fixed alleles

Minor allele frequency (MAF) and fixed allele proportion (i.e., MAF = 0) were obtained for each breed and SNP with the function *‘--freq’* in PLINK. Mean observed and expected heterozygosity were calculated using *‘--het’* in PLINK across all breeds and within each breed to compute the mean individual inbreeding coefficients based on allele frequencies, equivalent to F_IT a_nd F_IS,_ respectively (Wright, 1949).

#### Inbreeding and Kinship

POPKIN R package was used to compute pairwise kinship coefficients, as it accounts for arbitrarily complex population structure (Ochoa & Storey, 2021) as observed in our sample since we jointly considered all breeds. In addition, POPKIN makes estimates comparable across breeds by calculating Identity by Descent (IBD) probabilities on a scale relative to the Most Recent Common Ancestor (MRCA) which is helpful for making conservation decisions (Ochoa & Storey, 2021). The inbreeding coefficients were retrieved from the diagonal elements of the pairwise kinship matrix. Both global and cohort-level estimates of kinship and inbreeding coefficients were computed to assess genetic diversity across breeds.

Additionally, we used the RZooRoH R package (Bertrand et al., 2019; Druet & Gautier, 2017) to identify Homozygous-by-Descent (HBD) using the Hidden Markov Model (HMM) and obtain the individual autozygosity, i.e., realized inbreeding coefficients. The identified HBD segments were classified into five categories: below 2 Mb, 2-4 Mb, 4-8 Mb, 8-16 Mb, and greater than 16 Mb.

#### Statistical Comparisons

We statistically compared these diversity parameters (heterozygosity, F-statistics, mean MAF, and fixed alleles) between populations by fitting a mixed model for MAF (SNPs as a random effect) and a generalized linear model (GLM) for the proportions of fixed alleles, observed heterozygosity, and expected heterozygosity, assuming a binomial distribution. A linear model was fitted with breed as a fixed effect for F_IT a_nd F_IS._ Similarly, a linear model was also fitted to analyze the kinship and inbreeding coefficients with breed and cohort as fixed effects, including their interaction.

Pairwise least-squares means contrasts were estimated using the R package *emmeans* (Lenth & Piaskowski, 2025). The emmeans were used to compare significant differences between breeds and between the two cohorts within each breed, respectively. The statistical formulas are provided in Supplementary Tables S4 and S5. Additionally, we compared the distributions and average lengths of HBD segments between cohorts using the Kolmogorov-Smirnov test and Welch’s t-test, respectively.

#### Effective Population Size

NeEstimator V2 (Do et al., 2014) was used to calculate the contemporary effective population size. Using a single common input, the software calculated the effective population size using three methods based on different metrics: linkage disequilibrium (Waples & Do, 2008), heterozygote excess (Zhdanova & Pudovkin, 2008), and molecular coancestry (Nomura, 2008). Additionally, with the help of demographic information (number of breeding males and females), the sex-ratio-based effective population size (Wright, 1938) was calculated using the following formula:

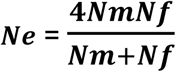

Where *Nm* and *Nf* are the numbers of breeding males and females, respectively. For the effective population size obtained from different measures, we also computed the ratio of effective population size against the actual breeding population, Ne/N. This allows evaluation of the management independently of the actual population size.

## 3. Results

### 3.1. Population Structure and Historical Diversity

The first axis of the Principal Component Analysis (PCA) explained 17% of the variance and showed a clear distinction between the draft and territory groups, with some breeds appearing intermediate, such as Cob Normand (COBND), Auvergne (AUV), Merens (MER), and Castillonnais (CAST). Axes 2 and 3 separated Boulonnais (BOUL) and Poitevin Mulassier (POIT), respectively, from the rest of the draft horses, while PC4 separated the breeds within the territory group (Figure 2).

**Figure 2:**
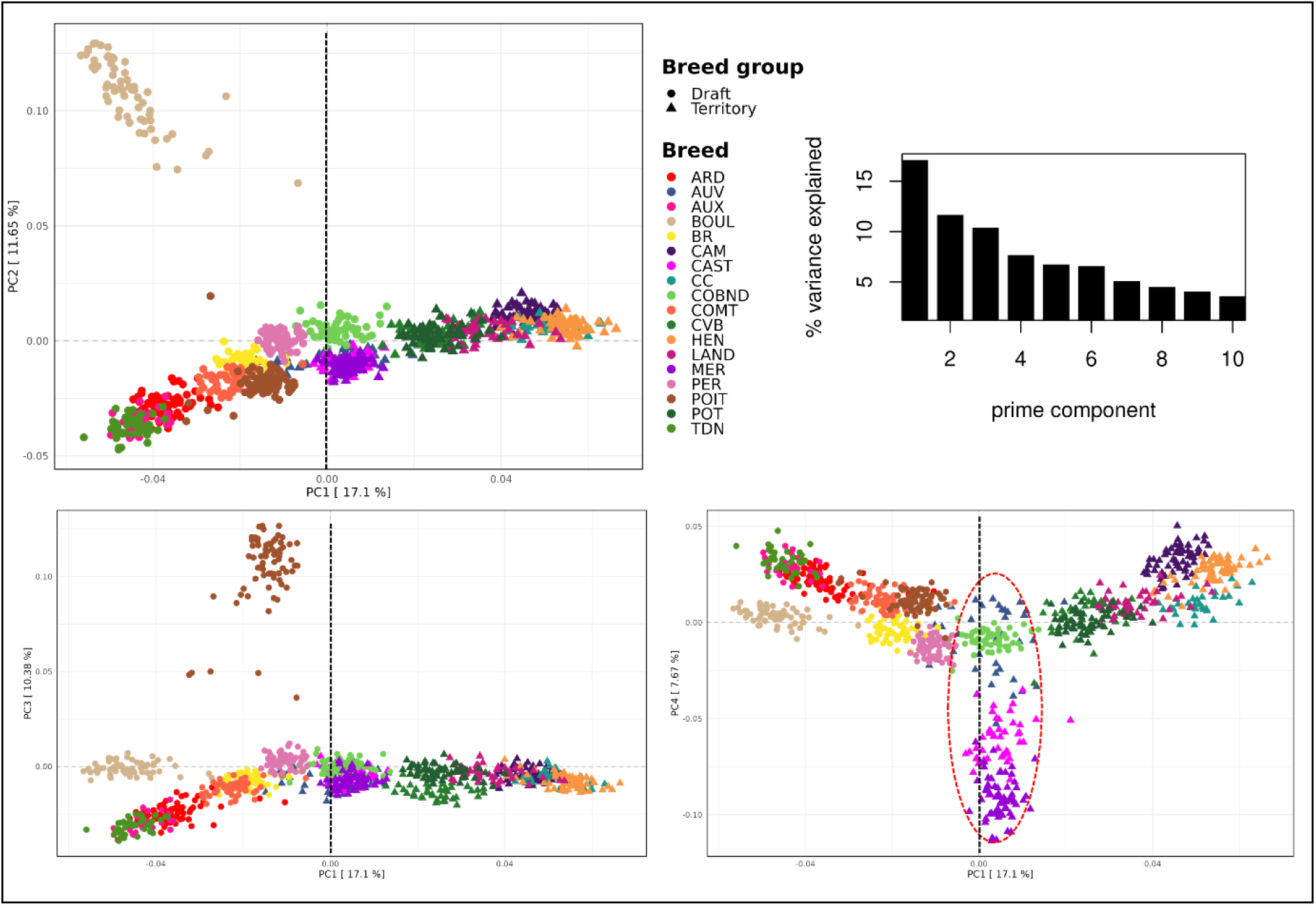
A) PCA with PC1 and PC2. B) Scree plot; C) PC2 versus PC3; D) PC3 versus PC4, the red dotted eclipse presenting intermediate breeds: COBND, AUV, MER, and CAST. The black dotted line represents the boundary between drafts and territory breeds. Breed abbreviations are given in Table 1.

For admixture analysis with a complete dataset, we did not identify any optimal *K* (i.e., the most likely number of coancestry groups) as the CV error values form a smooth curve without a noticeable drop (“elbow”). Therefore, we used a balanced dataset to identify the optimal *K*, which was found to be *K*=15 with the lowest CV error (0.611), as shown in Supplementary Figure S2. The correlation test between the ancestry allele frequency matrices at *K*=15 for both balanced and complete datasets showed a significantly high correlation of 0.86 (p-value < 2.2e-16) (Supplementary Figure 2), which allows us to interpret the optimal *K* in the latter dataset. At *K*=2, we observed a distinction between draft and territory breeds, with two dominant clusters forming a gradient from draft to territory, alongside intermediate breeds observed in PCA (Figure 3). At *K*=15, shared ancestry was observed among [Ardennais (ARD), Trait du Nord (TDN), Auxois (AUX)], [Pottok (POT), Camargue (CAM), Cheval Corse (CC)], [Auvergne, Cheval du Vercors de Barraquand (CVB/Vercors)], [Castillonnais (CAST), Merens (MER)], and [Castillonnais and Pottok]. Unlike at *K*=15, we observed dominant Cheval Corse and Pottok clusters at *K*=18. Furthermore, Pottok, Auvergne, and Vercors showed higher admixture than other breeds.

**Figure 3:**
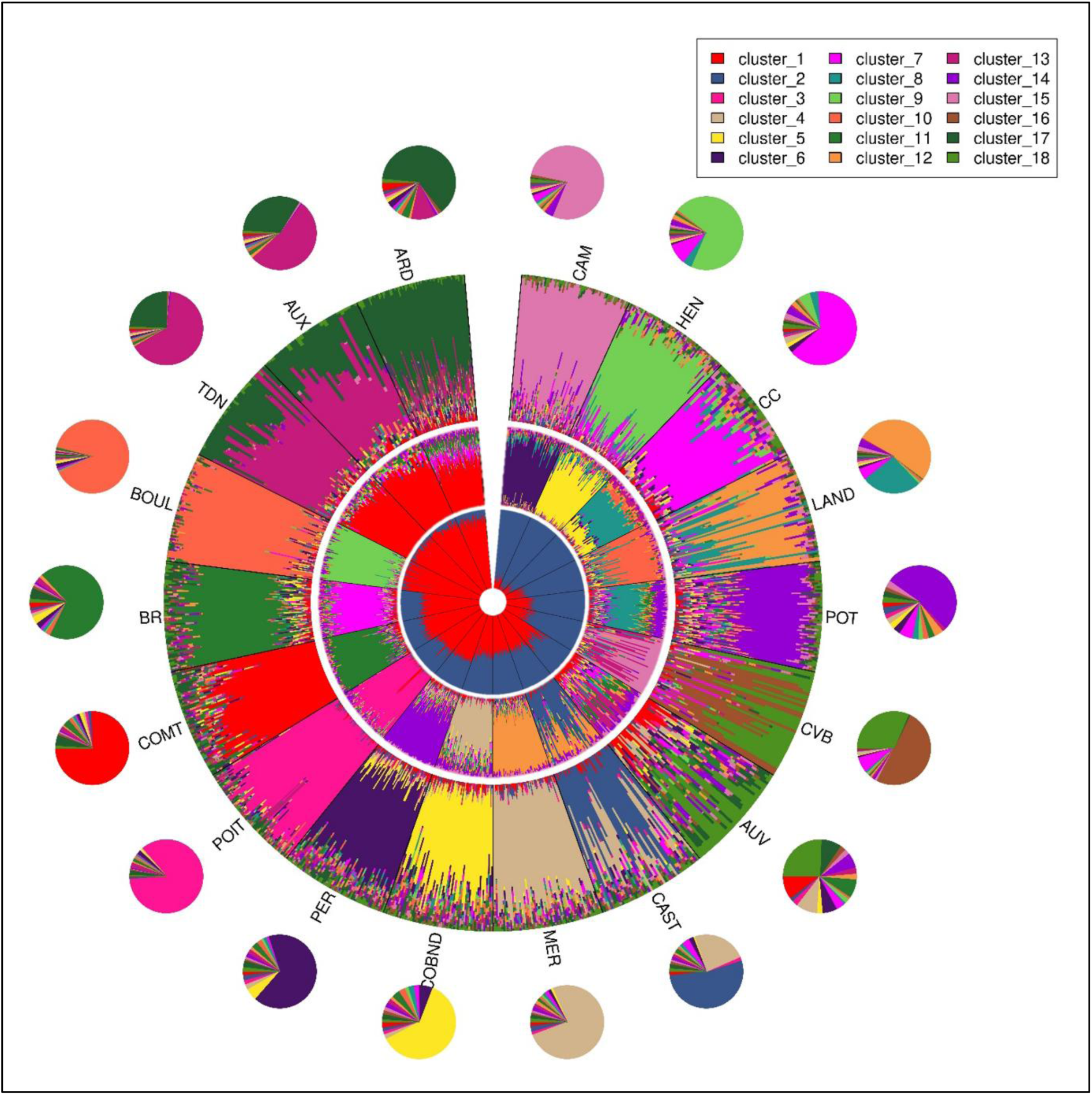
Admixture graphs of 18 horse breeds show K values of 2 (inner circle), 15 (middle circle), and 18 (outer circle), with average ancestry represented by a pie chart for the breeds.

The individual-based phylogeny (Supplementary Figure 3) supported the admixture analysis, with a single cluster comprising Auxois, Ardennais, and Trait du Nord, while most other individuals are well clustered within their respective breeds. Auvergne, one of the most admixed breeds, showed three clusters: one closer to the other mountain breeds, such as Merens, Castillonnais, and Vercors, and two closer to the draft breeds, such as Comtois.

Pairwise F_ST r_anged from 0.14 for Boulonnais vs Vercors to 0.008 for Trait du Nord vs Auxois (Supplementary Figure 4), with a global average of 0.072 (standard error: 0.002). An unrooted neighbour-joining tree based on the pairwise F_ST m_atrix showed a clear distinction between the two breed groups, with long branches for Boulonnais and Poitevin Mulassier (Figure 4). The rooted neighbour-joining tree based on the Reynolds distance showed that the territory breeds are closer to the ancestral population than the draft breeds, with Pottok appearing to be the root for most breeds, except for Cheval Corse, Henson, and Poney Landais. Moreover, the Treemix analyses showed a tree similar to the rooted tree, with two migration events added: from Comtois to Auvergne and from the roots of Cheval Corse, Henson, and Poney Landais to Cob Normand (Figure 4).

**Figure 4:**
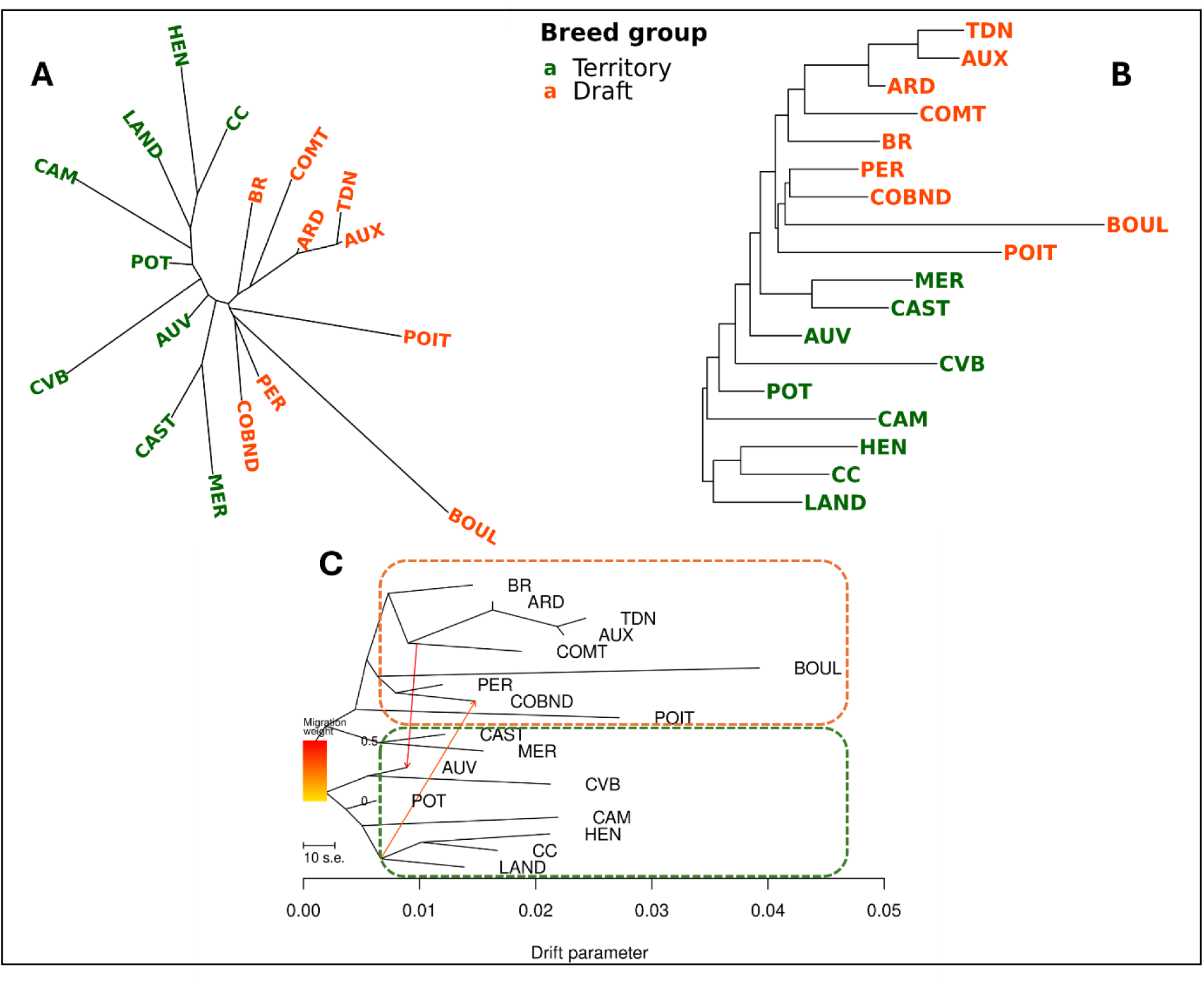
A) Unrooted neighbour-joining tree generated from FST data for 18 breeds, distinguished by two colours representing two breed groups. B) Rooted neighbour-joining tree created from the Reynolds distance, also using two colours to denote the two breed groups. C) Treemix graph illustrating two migration events, with migration arrows coloured according to their weight; darker arrows indicate heavier migration. The coloured dotted box highlights the two breed groups.

### 3.2. Landscape Genomics

The Mantel tests between the geographical and genetic distances were not significant for either draft or territory breeds (r = -0.0445, p = 0.587, and r = -0.513, p = 0.987, respectively) (Supplementary Figure S6). High correlations among the environmental variables were observed; hence, after backward variable selection, 13 variables (4 climatic, 5 soil, 3 land coverage, and elevation) were retained from the initial 30 variables (Supplementary Figure S5). We used *K* = 2 to account for breed-group structure (draft and territory), as observed in our population structure analyses, and retained 13 environmental variables for the final model. To account for partial redundancy, we performed variance partitioning as shown in Table 2, which revealed that the environmental variables explained 10.4% of the genomic variation after conditioning for the breeds’ group (adj. R^2^ = 0.092, p = 0.001). The axes 1 to 8 captured 84% of the cumulative variance explained by environmental factors, such as sand, soil pH_, v_olume fraction of coarse fragments, barren land, cropland, and elevation, with a correlation with RDA axes greater than 0.5 (Supplementary Figure S5).

**Table 2:** The influence of environment and population structure on genetic variation decomposed with pRDA (partial redundancy analysis)

| RDA models | Inertia | R2 | p (>F) | Proportion of explainable variance | Proportion of total variance |
| --- | --- | --- | --- | --- | --- |
| Full model | 360.1 | 0.115 | 0.001 *** | 1 | 0.128 |
| Environment | 291.5 | 0.092 | 0.001 *** | 0.81 | 0.104 |
| Structure<br>(Draft vs territory) | 24.66 | 0.008 | 0.001 *** | 0.07 | 0.009 |
| Confounded<br>(Environment/structure) | 43.86 |  |  | 0.12 | 0.016 |
| Total unexplained | 2455.2 |  |  |  | 0.87 |
| Total inertia | 2815.2 |  |  |  | 1 |

### 3.3. Genetic Diversity and Management

MAF, the fixed-allele proportion, expected heterozygosity, observed heterozygosity, F_IS,_ and F_IT a_re summarized in Table 3. The minor allele frequency ranged from 0.26 (Boulonnais) to 0.34 (Pottok), with a global mean of 0.31. Pottok had the significantly highest MAF and the lowest fixed allele proportion (0.014) compared to all the other breeds, while Boulonnais ranked as the breed with the highest fixed allele proportion (0.05). Expected heterozygosity ranged from 0.36 for Boulonnais to 0.42 for Pottok, values close to the observed heterozygosity, which ranged from 0.37 for Boulonnais to 0.43 for CC, with a global average of 0.4 and 0.41, respectively. The average F_IS f_or all breeds is negative, except for Camargue (0.008), which differs significantly from the other breeds. On the other hand, the average F_IT w_as positive and followed the pattern of MAF and fixed-allele proportion, with Pottok having the lowest value (0.0058) and Boulonnais the highest (0.167).

**Table 3:** Summary statistics of diversity parameters.

| Breed | MAF | Fixed allele | Exp. Heterozygosity |  |  | Obs. Heterozygosity |  |  | F<sub>IS</sub> |  |  | F<sub>IT</sub> |  |  |
| --- | --- | --- | --- | --- | --- | --- | --- | --- | --- | --- | --- | --- | --- | --- |
|  | Mean | Proportion | Mean | Min | Max | Mean | Min | Max | Mean | Min | Max | Mean | Min | Max |
| ARD | 0.306<sup>de</sup> | 0.037<sup>abc</sup> | 0.399<sup>def</sup> | 0.398 | 0.399 | 0.405<sup>fg</sup> | 0.372 | 0.59 | -0.016<sup>bc</sup> | -0.481 | 0.068 | 0.067<sup>cd</sup> | -0.364 | 0.138 |
| AUV | 0.327<sup>h</sup> | 0.044<sup>bcdef</sup> | 0.413<sup>gh</sup> | 0.412 | 0.413 | 0.424<sup>lm</sup> | 0.355 | 0.448 | -0.028<sup>abc</sup> | -0.086 | 0.141 | 0.01<sup>a</sup> | -0.044 | 0.169 |
| AUX | 0.292<sup>b</sup> | 0.074<sup>h</sup> | 0.39<sup>c</sup> | 0.389 | 0.39 | 0.403<sup>defg</sup> | 0.355 | 0.424 | -0.034<sup>abc</sup> | -0.087 | 0.087 | 0.09<sup>def</sup> | 0.036 | 0.188 |
| BOUL | 0.262<sup>a</sup> | 0.058<sup>g</sup> | 0.359<sup>a</sup> | 0.359 | 0.359 | 0.369<sup>a</sup> | 0.313 | 0.432 | -0.029<sup>abc</sup> | -0.203 | 0.128 | 0.168<sup>g</sup> | 0.018 | 0.292 |
| BR | 0.308<sup>de</sup> | 0.04<sup>abcd</sup> | 0.401<sup>f</sup> | 0.401 | 0.402 | 0.41<sup>h</sup> | 0.374 | 0.433 | -0.022<sup>abc</sup> | -0.08 | 0.069 | 0.055<sup>cd</sup> | -0.001 | 0.14 |
| CAM | 0.306<sup>de</sup> | 0.043<sup>bcdef</sup> | 0.399<sup>def</sup> | 0.398 | 0.4 | 0.399<sup>cd</sup> | 0.314 | 0.438 | 0.002<sup>c</sup> | -0.097 | 0.214 | 0.082<sup>def</sup> | -0.007 | 0.275 |
| CAST | 0.311<sup>e</sup> | 0.048<sup>cdefg</sup> | 0.4<sup>ef</sup> | 0.4 | 0.4 | 0.412<sup>hi</sup> | 0.324 | 0.436 | -0.03<sup>abc</sup> | -0.091 | 0.189 | 0.044<sup>bc</sup> | -0.022 | 0.252 |
| CC | 0.325<sup>gh</sup> | 0.055<sup>fg</sup> | 0.413<sup>gh</sup> | 0.411 | 0.413 | 0.428<sup>m</sup> | 0.398 | 0.491 | -0.035<sup>abc</sup> | -0.194 | 0.038 | 0.008<sup>ab</sup> | -0.147 | 0.084 |
| COBND | 0.324<sup>gh</sup> | 0.039<sup>abcd</sup> | 0.413<sup>h</sup> | 0.413 | 0.414 | 0.423<sup>kl</sup> | 0.394 | 0.491 | -0.022<sup>abc</sup> | -0.19 | 0.047 | 0.021<sup>ab</sup> | -0.14 | 0.091 |
| COMT | 0.303<sup>d</sup> | 0.049<sup>cefg</sup> | 0.397<sup>de</sup> | 0.397 | 0.397 | 0.4<sup>de</sup> | 0.357 | 0.437 | -0.007<sup>bc</sup> | -0.101 | 0.101 | 0.084<sup>def</sup> | -0.009 | 0.177 |
| CVB | 0.297<sup>c</sup> | 0.053<sup>efg</sup> | 0.388<sup>c</sup> | 0.388 | 0.389 | 0.409<sup>gh</sup> | 0.317 | 0.464 | -0.053<sup>a</sup> | -0.196 | 0.183 | 0.063<sup>cd</sup> | -0.065 | 0.285 |
| HEN | 0.32<sup>fg</sup> | 0.036<sup>ab</sup> | 0.409<sup>g</sup> | 0.408 | 0.409 | 0.424<sup>lm</sup> | 0.369 | 0.454 | -0.038<sup>ab</sup> | -0.114 | 0.097 | 0.014<sup>ab</sup> | -0.057 | 0.137 |
| LAND | 0.326<sup>h</sup> | 0.041<sup>abcde</sup> | 0.411<sup>gh</sup> | 0.411 | 0.411 | 0.419<sup>jk</sup> | 0.372 | 0.448 | -0.018<sup>abc</sup> | -0.089 | 0.096 | 0.023<sup>ab</sup> | -0.047 | 0.133 |
| MER | 0.304<sup>d</sup> | 0.041<sup>abcd</sup> | 0.396<sup>d</sup> | 0.394 | 0.396 | 0.403<sup>ef</sup> | 0.37 | 0.44 | -0.018<sup>abc</sup> | -0.117 | 0.065 | 0.07<sup>cde</sup> | -0.024 | 0.152 |
| PER | 0.318<sup>f</sup> | 0.039<sup>abcd</sup> | 0.409<sup>gh</sup> | 0.408 | 0.41 | 0.415<sup>ij</sup> | 0.363 | 0.473 | -0.014<sup>bc</sup> | -0.16 | 0.114 | 0.042<sup>bc</sup> | -0.093 | 0.162 |
| POIT | 0.288<sup>b</sup> | 0.046<sup>bcdefg</sup> | 0.382<sup>b</sup> | 0.382 | 0.383 | 0.391<sup>b</sup> | 0.329 | 0.438 | -0.023<sup>abc</sup> | -0.146 | 0.14 | 0.103<sup>f</sup> | 0.006 | 0.244 |
| POT | 0.342<sup>i</sup> | 0.032<sup>a</sup> | 0.423<sup>i</sup> | 0.423 | 0.424 | 0.425<sup>lm</sup> | 0.341 | 0.448 | -0.003<sup>c</sup> | -0.058 | 0.195 | 0.006<sup>a</sup> | -0.047 | 0.206 |
| TDN | 0.29<sup>b</sup> | 0.057<sup>g</sup> | 0.386<sup>bc</sup> | 0.385 | 0.386 | 0.395<sup>bc</sup> | 0.368 | 0.423 | -0.024<sup>abc</sup> | -0.097 | 0.046 | 0.101<sup>ef</sup> | 0.05 | 0.163 |
| Mean | 0.308 | 0.046 | 0.399 | 0.399 | 0.4 | 0.409 | 0.355 | 0.456 | -0.023 | -0.144 | 0.112 | 0.058 | -0.053 | 0.183 |

The letters represent significant differences in the diversity parameter when compared across the breeds, with letter “a” denoting the smallest value.

The same trend was observed for inbreeding and kinship coefficients at the global level (Table 4), with Pottok having a significantly lower kinship coefficient of 0.037 and the lowest inbreeding of 0.043, while Boulonnais had the highest values for both estimates, 0.215 and 0.199, respectively.

**Table 4:**
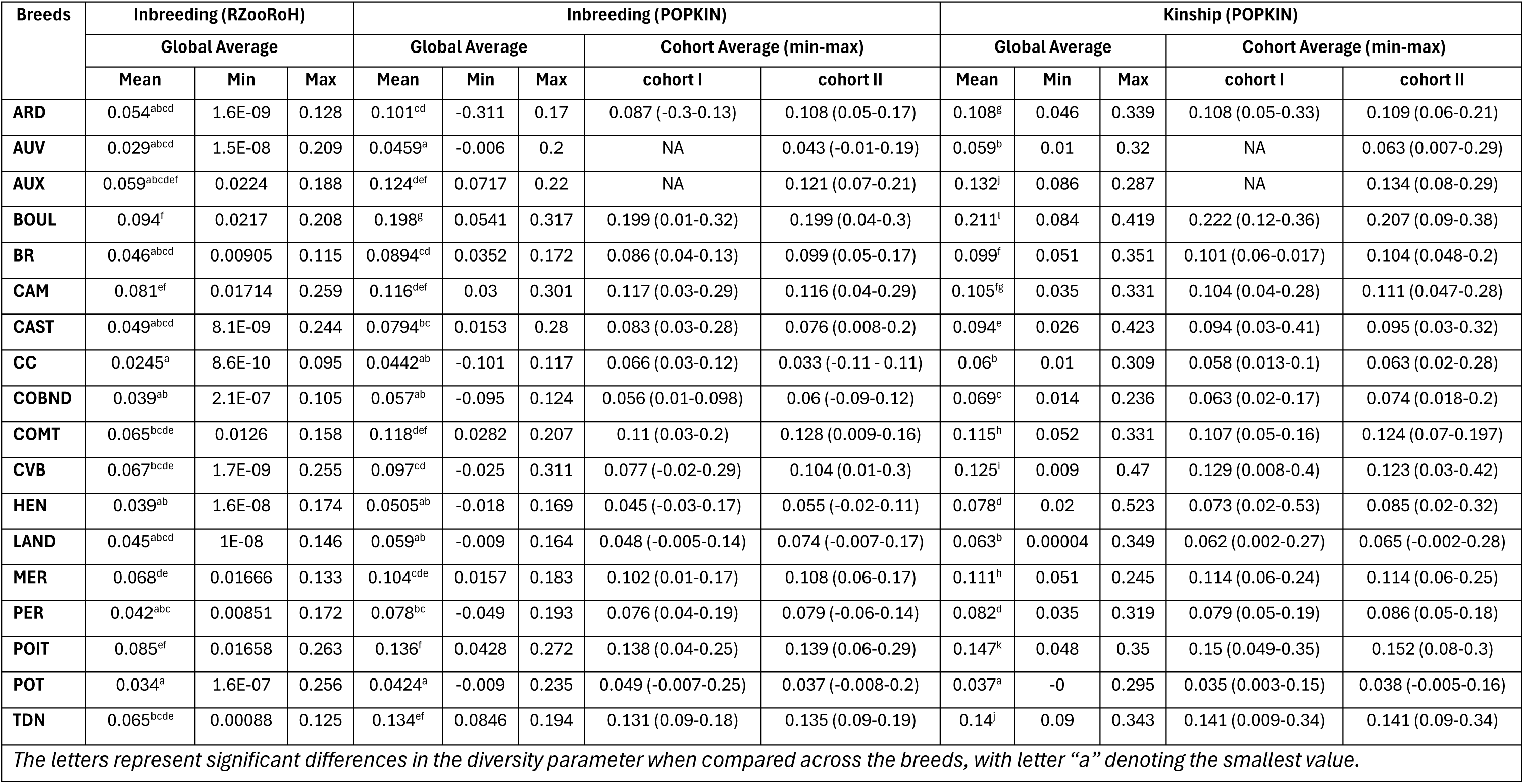
Summary of Inbreeding and kinship coefficients obtained from the R packages: POPKIN and RZooRoH.

Nevertheless, Vercors, with kinship (0.13) and inbreeding coefficients (0.09) close to the global means (0.104 and 0.903, respectively), had high variability in kinship (0.03 to 0.42) and inbreeding coefficients (0.01 to 0.3). At the group level, the average inbreeding coefficients of territory and draft groups are 0.07±0.002 versus 0.11±0.002, respectively. Among the ten breeds with inbreeding coefficients of 10% or higher, eight are draft horses. At the global level, F_IT h_ad perfect Spearman’s rank correlation (p-value: 1.4e-47) with inbreeding coefficients obtained from POPKIN and 0.86 with RZooRoH estimates (p-value: 7.7e-08), while F_IS h_ad a higher correlation with RZooRoH (0.72, p-value: 3.1e-283) than POPKIN (0.65, p-value: 1e-145).

Looking at the length of HBD segments, all breeds had a higher number of HBD segments between 4 and 8 Mb (35% of the total HBD detected), followed by 8 and 16 Mb (32%) (Supplementary Table S7). Ardennais has the lowest average length of HBD segments detected (7.5 Mb), and Vercors has the highest (16.29 Mb), with a wide range (2.05 Mb–98.2 Mb), following Camargue (1.5 Mb-100.67 Mb) (Supplementary Table S8). Moreover, Vercors has the highest mean individual HBD coverage (172.3 Mb), while Cheval Corse has the least (42.64 Mb). The highest mean number of HBD segments per individual is observed in Camargue (13.24) and the lowest in Cheval Corse (4.07). The Kolmogorov-Smirnov test shows that the distribution of HBD segment lengths differs significantly between the two cohorts for Camargue and Poney Landais. Additionally, Welch’s t-test indicates a significant difference in the average HBD segment length between the Breton and Castillonnais cohorts.

Moreover, we assessed the evolution of inbreeding and kinship coefficients within breeds between cohorts (Figure 5). We observed a general trend of increasing inbreeding with increasing kinship at the global level; however, at the breed level, inbreeding and kinship evolved differently, as shown by arrows pointing in different directions. Boulonnais showed a significant decrease in the kinship coefficient, whereas Camargue, Cob Normand, Henson, Comtois, and Percheron showed a significant increase. When we examined the inbreeding coefficient, only Cheval Corse decreased significantly, whereas Vercors and Poney Landais increased significantly. The emmeans differences between estimates from cohorts I and II broadly clustered the breeds into four groups; some breeds fell between two groups, leading to six categories (Table 5). At the cohort level, the Spearman’s rank correlation between F_IT a_nd inbreeding estimates from POPKIN is 0.97(p-value: < 2.2e-16), whereas for F_IS i_t is 0.35 (p-value: < 2.2e-16).

**Table 5:** Evolution of Inbreeding and Kinship coefficients across two cohorts Category (Cohort I ◊ II) Breeds.

| Category (Cohort I → II) | Breeds |
| --- | --- |
| Higher inbreeding and kinship | POIT, COMT, BR, PER, HEN, COBND, LAND, ARD |
| Lower inbreeding, higher kinship | CAST, CC, POT |
| Higher inbreeding and lower kinship | CVB |
| No change in inbreeding, higher kinship | CAM |
| No change in inbreeding, lower kinship | BOUL |
| No change in kinship, higher inbreeding | TDN, MER |

**Figure 5:**
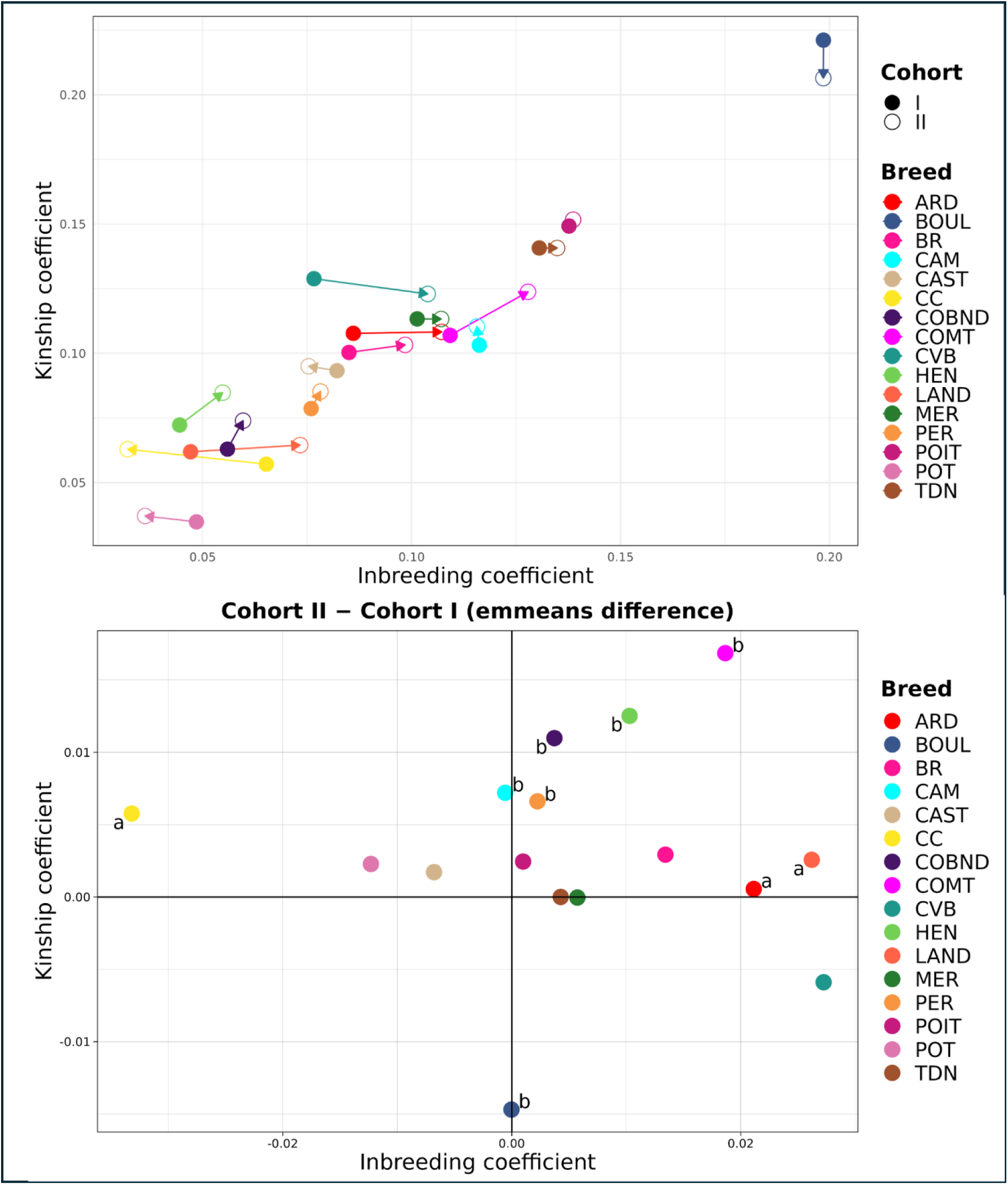
The y-axis represents the kinship coefficient, and the x-axis represents the inbreeding coefficient. A) The full circle presents estimated values for kinship and inbreeding coefficients for cohort I, and the empty circle for cohort II, with the arrow moving from cohort I to II. B) The points present differences in the estimated values between cohort II and cohort I, with subscript ‘a’ denoting significant differences in the inbreeding coefficient, and ‘b’ denoting significant differences in the kinship coefficient. The highlighted axis presents lines at x=0 and y=0, which divide the graph into four quadrants.

The effective population sizes for each method showed estimates with varying trends and values (Figure 6). The average value for Demographic is the highest at 406.4, followed by Linkage Disequilibrium at 166.7, Heterozygous Excess at 32.07, and Molecular Coancestry at 9.88. We observed several instances of infinite values with the heterozygous excess method. The ratio of effective population size to the total breeding population presents a different picture than effective population size alone, with Cheval Corse showing the highest ratio and lower ratios for breeds with higher Ne, such as Ardennais, Breton, Comtois, and Percheron (Supplementary Table S6).

**Figure 6:**
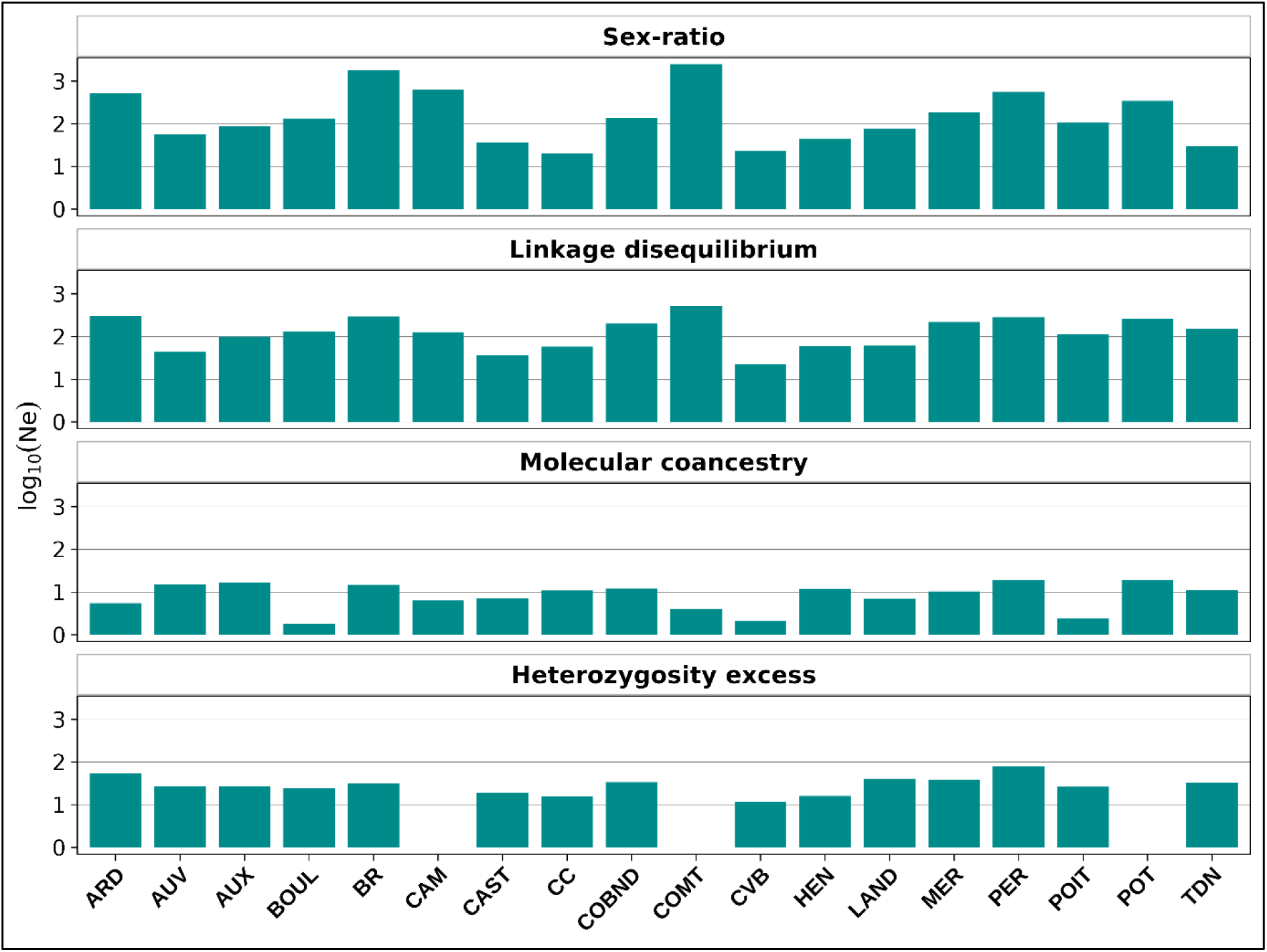
Log10-transformation of Effective Population Size (Ne) obtained from four different methods, namely, sex-ratio, Linkage Disequilibrium, Molecular coancestry, and Heterozygosity excess. The gaps in the heterozygosity excess method are due to infinite values.

## 4. Discussion

### History of the local breeds

Using several methods, we reconstructed both the past history of all recognized French local horse breeds and their fates through the evolution of genetic diversity.

#### Breed formation

French local horses can be classified not only phenotypically and from a “cultural” point of view but also genetically into two categories: draft and territory groups, as validated by PCA, admixture analysis, and all the independent tree-based analyses (Figures 1, 2 & 3). Among these two distinct groups, territory breeds (except Cheval Corse, Henson, and Poney Landais) form the basal lineage to draft breeds and appear closer to the ancestral horses (Figure 4). This phylogenetic order is plausible, as draft breeds were developed later in the 18th and 19th centuries in response to increased demand for heavy horses for agricultural and heavy work (Gosselin, 2023b). Further, the breeds forming the root, like Pottok and Camargue, can be traced back to the wild horses of the Basque Country, dating to the Paleolithic era (Paulat-Boucher, 2010; Rousseaux, n.d.) and the Iberian, and even Solutré horses (Rousseaux, n.d.) respectively.

Further, we observed intriguing structures and relationships among the breeds within the groups. In the draft group, the most notable finding was the shared ancestry among the Ardennais, Trait du Nord, and Auxois breeds, which was supported by both admixture and tree-based analyses in line with previous studies (Leroy et al., 2009). The Ardennais is recognized as one of the oldest draft breeds in Europe, known for its role as a breed improver and its contribution to the development of other draft breeds such as Trait du Nord and Auxois, which helps explain the observed shared ancestry among these breeds (FCC, 2022b). In contrast, we noted that Boulonnais and Poitevin Mulassier diverged, as evidenced by PCA and the longest branches in the phylogenetic trees, particularly for Boulonnais (refer to Figures 1 & 3). Our analyses using various diversity parameters further also supported this strong drift. Historical records regarding Boulonnais management reveal strong selection pressure on coat colour; until the 18th century, these animals predominantly had black and dark grey coats. However, a preference developed for light grey coats, as this coloration made it easier to see the horses while transporting fish carts at night, ultimately leading to the selection for this specific coat colour (EQWOS, 2016). On the other hand, Poitevin Mulassier were selected for their ability to produce Poitevin mules, which lost their popularity in the middle of the 20th century after mechanization, leading this breed to decline in population size and to suffer from strong inbreeding, also because half of its offspring were infertile mules rather than the horses themselves (Trait Poitevin Mulassier(Cheval), 2021). According to a previous study on genetic diversity and structure among French horse breeds using microsatellite markers, Boulonnais, despite diverging from the other breeds, has retained genes from its original gene pool (Leroy et al., 2009). Conversely, Poitevin Mulassier, while exhibiting the least allelic diversity, contributed significantly to between-breed diversity alongside Boulonnais (Leroy et al., 2009). This underscores the uniqueness of these breeds and highlights their importance for conservation efforts.

Within territory groups, relatedness among the breeds can be traced back in older time than in draft horses. For instance, the shared ancestry between Pottok and Cheval Corse (*K*=15) is believed to result from the introduction of Oriental bloodlines (Arabian or Barb) during the Saracen invasion in the 8th century (Rousseaux, n.d.; U Cavallu Corsu, n.d.). Further, the prevailing hypothesis supporting the closeness between Merens and Castillonnais is that the ancestors of these breeds may have introgressed with wild Ariège horses, as these breeds share remarkable morphological similarities with Magdalenian horses (13000 years ago) depicted in the Ariège caves (Horse Isle, n.d.; Rousseaux, n.d.). However, over time, Castillonnais was influenced by Oriental (Arabs and Barbs) and the Spanish (Iberian) breeds (Horse Isle, n.d.; Rousseaux, n.d.). Hence, an alternative and maybe more realistic explanation of their relatedness could be their shared geographic proximity and a similar mountainous climatic zone (SFET, n.d.-a). Similarly, the shared ancestry between Vercors and Auvergne is not straightforward, but, like the previous breeds, they are geographically close and thrive in mountainous regions (SFET, n.d.-b). Moreover, this could also be because Vercors experienced several bottlenecks in the past, most significantly after the heavy losses in the Second World War, and was revived with a handful of remaining mares and stallions; in that process, Auvergne could have been used to recover the population.

The small recovery population with some out-crossing could explain the family-like substructure of Vercors in admixture analyses. Finally, the Poney Landais, Cheval Corse, and Henson form a separate branch in the rooted tree, which could result from a significant amount of cross-breeding these breeds had undergone with Arabian horses for improvement (Horse Isle, n.d.).

Along with relationships within groups, we also observed relatedness between the groups either through gene flow or selection in response to the needs of the time, which has resulted in some breeds (Auvergne, Castillonnais, Cob Normand, and Merens) appearing as intermediates (PCA: PC1 axis and admixture proportions at K=2). The Auvergne is believed to have originated from Oriental breeds left behind by the Saracen army after its defeat in the 8th century and later developed into a distinct breed by the 12th century, roaming the mountains of Auvergne. In the 17th century, this breed split into two distinct groups: one through cross-breeding with English and Anglo-Arab horses, used as a war horse, and a second branch of primitive horses used for farming. But by the end of the 18th and 19th centuries, the primitive types were heavily crossed with draft breeds like Comtois, which we observed in our Treemix analysis (Figure 4C) (FCC, 2022a; Horse Isle, n.d.). Similarly, Castillonnais, with a high demand for heavy horses, was largely crossbred with draft horses to the point that it became almost extinct when mechanization began (Horse Isle, n.d.). While these territory breeds were crossed with heavy horses for agricultural work, Cob Normand, a draft horse, was crossed with Norfolk trotter in the 19^th^ century and was promoted as a light draft (Horse Isle, n.d.; Rousseaux, n.d.). Lastly, unlike the horses above, where a new bloodline was introduced, Merens (a small mountain horse) was selected for heavier size to perform heavy agricultural work and is also called a light draft horse (FCC, n.d.).

Beyond breed relatedness, we found interesting insights from admixture analysis, with some breeds appearing highly admixed at K=18. Unusually, Pottok, being an ancient breed, appeared heterogeneous, which could be because the Admixture model assumes that individuals receive contributions from one or more of K genetic ancestral clusters (Pritchard et al., 2000) and in some cases, the model can assign individual clusters to the derived population while assigning high admixture proportions to the ancestral population in an attempt to fit the data as best it can (Lawson et al., 2018). Moreover, Auvergne appeared highly admixed, which could be due to the presence of different types, one closer to the original and the other with multiple foreign gene flows, as revealed in the individual identity-by-state (IBS) neighbor-joining tree with multiple clusters (Supplementary Figure S3). In contrast, Henson, a newly formed breed by crossing first-generation offspring of a Fjord pony and a saddle horse, did not appear as admixed, which could be due to a closed breeding among the first-generation offspring since 1955, creating a single nucleus for a breed to develop, resulting in a genetically stabilized population (SFET, n.d.-a).

#### Local adaptation of breeds

First, we tested isolation by distance that can blur the signal for local adaptation, and found no significant correlation between genetic and geographic distances for each breed group. This could be due to a mainly human-driven structure inducing limited or directed gene flow (i.e., breed structure), or to adaptation to local environments. Isolation by distance was tested separately for each breed group (i.e., draft or territory) to avoid upward bias, as draft horses are concentrated in the Northern region of France, whereas most territory horses are in the Southern regions.

We conducted a variable selection step to account for collinearity, which could lead to variance inflation and biased predictor identification (Dormann et al., 2013) and obtained a final set of 13 variables, including climatic, soil, land cover, and elevation variables. Based on our previous observation, we used the strongest source of genetic structuring (*K*=2) as a covariate, allowing us to focus on major population clustering (draft versus territory groups) while retaining enough power to detect signals of local adaptation within groups. Moreover, the local breeds in our study are sampled from one single geographic region, which is a typical feature of traditional livestock breeds, lacking spatial/environmental replication unlike commercial breeds and plant species, making classical landscape genomics more challenging (Li et al., 2017; Richardson et al., 2016).

The environmental variables explained more variation than the group effect (draft and territory), when conditioned for one another. However, we found a confounding effect that suggests collinearity between the predictors (environmental and structural), such that nearly 12% of the explanatory variation cannot be associated with any of the predictors (Capblancq & Forester, 2021). Environmental factors associated with land coverage, soil, and elevation seemed to exert a stronger influence than climatic factors (Supplementary Figure 5), which may be closely tied to the horses’ purpose rather than to the environment itself. For instance, Camargue, also known as “the horse of the sea,” is used for agricultural purposes on the swampy beaches of the Camargue, which, in the RDA bi-plot, is clustered with cropland (Supplementary Figure 5). Similarly, mountain horses such as Merens and Castillonnais are used for hiking in difficult terrain because they are known for their sure-footedness, and we observed these breeds clustering with high elevation (Supplementary Figure 5). Our analyses at this point cannot distinguish whether adaptation is related to environment or purposes of these horses; however, it provides a strong base for future studies with a dataset having finer-scale environmental variations and high SNP density to be able to detect specific adaptive genetic loci associated with environment or selection signatures, as in some prevalent studies in other species (Capblancq & Forester, 2021; Vallejo-Trujillo et al., 2022; Senczuk et al., 2022; Zhang et al., 2023; Di Civita et al., 2026).

### Recent genetic evolution of local breeds

While French local breeds are numerous and have experienced contrasted histories leading to such a large diversity of phenotypes locally adapted to both environment and practices, their conservation is now at risk since all of them are threatened (Allier et al., 2023; Verrier et al., 2015; 2023). Thus, it is necessary to evaluate their genetic diversity and their management to conserve these valuable genetic resources.

#### Evaluation of genetic diversity and conservation status

We observed negative F_IS v_alues in most of our breeds, indicating higher heterozygosity than expected, except for Camargue (0.008). Higher expected heterozygosity can occur for several reasons, such as finite populations, which is the case for endangered breeds as in our study (Waples, 2015). Finite population size can lead to chance differences in genotype frequencies between male and female parents, resulting in excess heterozygosity relative to expectations in the progeny (Robertson, 1965). This could be the case for Cheval Corse because of its small breeding population (19 mares and 7 stallions). It may also reflect an isolate-breaking effect from recent outcrossing among isolate founders, as in the newly formed breed, Henson. Also, selection differences in male and female populations and strong avoidance of mating with close relatives could result in higher heterozygosity (Waples, 2015; Wright, 1965). The last scenario presents a best-case in terms of management, and France has implemented an online tool known as Cupidon that helps breeders access information on reproductive animals (mares and stallions), providing personalized advice encouraging breeders to conserve genetic diversity (Bonnin et al., 2019). Lastly, SNP ascertainment bias could lead to a systematic overestimation of heterozygosity in underrepresented breeds in SNP arrays (Clark et al., 2005; Geibel et al., 2021). On the other hand, positive F_IS o_r lower-than-expected heterozygosity can happen in cases of the Wahlund effect (population substructure), null alleles, and positive assortative mating (Waples, 2015; Wright, 1965). To rule out the temporal Wahlund effect, as we have samples from two cohorts, we looked at the F_ST b_etween the breeds across the cohorts and observed it to be negligible, ranging from -0.008 for Poney Landais to 0.004 for BR.

While F_IS i_ndicates the effectiveness of population management, given the allelic diversity within the population, F_IT i_ndicates the diversity retained in the sub-populations after the common ancestral pool diverged (Muir et al., 2008; Waples, 2015; Restoux et al., 2022). Since F_IS p_rovides information on the most recent generation of mating, it does not provide a straightforward summary of inbreeding; on the other hand, F_IT s_hows a higher correlation with inbreeding coefficients from POPKIN (1, p-value: 1.4e-47) and RZooRoH (0.86, p-value: 7.7e-08) and better reflects overall homozygosity and loss in diversity at the meta-population level. In Petersen et al. (2013), the highest inbreeding coefficient calculated from observed and expected heterozygosity at the total population level (F_IT)_ was found in the Mangalarga Paulista, a riding horse from Brazil (0.242), followed by the highest individual value of 0.56 in the Exmoor pony from Great Britain. The highest individual inbreeding coefficient in our study is 0.32 (Boulonnais), which is still lower than that reported in the previously mentioned study. The inbreeding coefficients of French horses are comparable to those of native horse breeds in Xinjiang (Tang et al., 2025) despite their expected heterozygosity being lower than in our study. Finnish and Iranian horses have a much lower inbreeding coefficients of 0.032 (Kvist et al., 2019) and 0.027 (Bazvand et al., 2024), respectively.

The higher inbreeding in draft horses (0.11±0.002, mean±s.e.) versus territory (0.07±0.002, mean±s.e.) may be because they were strongly selected for their build, but after the development of motors, horse-driven work declined, leading to a drastic decline in their population (Cressent & Jez, 2013). While the utility of territory horses is gaining popularity for alternative uses like leisure, draft horses are used less for this purpose. To still conserve these animals, their use shifted toward meat production (mainly abroad), and they were selected for it. This suggests that draft horses have undergone multiple rounds of strong, human-driven selection, along with the risk of population decline due to limited usefulness.

Further analysis of the HBD segments allows determination of the age of inbreeding, enabling inference of whether it is due to past history or recent events. According to the studies by Mulim et al. (2022), ROH lengths up to approximately 6 Mb are considered ancient, 6-12 Mb moderate, and lengths above 12 Mb recent inbreeding. In addition, Cardoso et al. (2020) referred to ROH of 8 Mb as formed more than 6 generations ago. Given that most HBD in our study (64.7%) are under 16 Mb, we can consider that inbreeding is fairly moderate in timeline. However, 19% of HBD segments in our study are over 16 Mb, with the longest HBD being 100.6 Mb (Camargue), which raises concerns about recent management practices. However, the large size of HBD in our study could be due to the low density and limited coverage of the SNPs, since low-to-moderate density SNP arrays may overestimate HBD segment lengths due to the absence of non-HBD SNPs flanking the segments (Solé et al., 2017). However, unlike traditional sliding-window methods that rely on fixed thresholds like Plink software (Purcell et al., 2007; Chang et al., 2015), the detection of HBD segments with a Hidden Markov Model (HMM) provided a robust probabilistic framework for accurately detecting real autozygosity.

A combination and comparison of various diversity parameters are essential for understanding a population’s genetic diversity and its evolution. Regardless of the parameters considered (Table 3), Boulonnais from the draft group appears to be the least genetically diverse and most drifted among the breeds, a pattern also observed in the assessment of population structure (PCA and phylogenetic trees) and supported by management history. It has the highest inbreeding and kinship estimates across breeds (0.199 and 0.213, respectively). In contrast, Pottok from the territory group appears to be the most genetically diverse, with the lowest values of inbreeding and kinship coefficients (0.042 and 0.037, respectively). The rich genetic diversity of Pottok was highlighted by Leroy et al. (2009), who reported the highest observed and expected heterozygosity, mean number of alleles, and allelic richness among the 13 breeds common to our study.

Effective population size predicts the loss and distribution of genetic diversity, which makes it a crucial parameter in conservation biology, yet one of the most challenging to estimate (Wang, 2005). In this study, we observed varying trends and values across four different approaches (Figure 6, Supplementary Table S6). These discrepancies arise from the different calculation methods, assumptions, and timescales employed in each of the metrics (Nomura, 2008; Waples & Do, 2008; Zhdanova & Pudovkin, 2008). We observed larger values for the sex-ratio and LD methods, as these methods estimate contemporary effective population size rather than long-term historical average. This may also stem from failing to account for relatedness among individuals, unlike molecular coancestry. That said, the lower values for molecular coancestry could reflect its capture of long-term drift, as seen for Boulonnais and Poitevin Mulassier (1.8 and 2.4, respectively). It was observed that this method introduces a downward bias when samples are drawn from an inbred or structured population (Nomura, 2008), which could have led to lower values, such as 2.1 for Vercors. Moreover, we obtained infinite values for three breeds (Camargue, Comtois, and Pottok) using the heterozygous excess method, which may be due to increased sampling error. Do et al. (2014) also observed infinite values, with 30% of results being infinite.

When comparing the study on effective population size of Finnish horses with our study, we observed that local French horses have a greater range of Ne (44 for Auvergne to 524.9 for Comtois) across breeds compared to Finnish horses (ranging from 43 for pony-sized horses to 56 for draught-type), but a similar value at the country level, with 166.7 (average across all French breeds) versus 161 (computed with Finnish breeds together), respectively. We observed a higher values of Ne for landraces from Mongolia (743), Tuva (533), and Iceland (555), as reported by Petersen et al. (2013). However, the Ne estimates in the study are obtained by combining all breed types within a specific country, rather than averaging them, which could have inflated them relative to our average Ne for French local horses (Petersen et al., 2013).

We also computed the ratio of effective population size to breeding population (Supplementary Table S6) and observed contrasting results for Comtois, with a value of 0.08 versus 0.4 for Auvergne. This highlights that the Ne/N ratio better reflects the potential genetic diversity given population size, providing important information for breed management and conservation prioritization (Jean Uzans, 2013; Waples, 2024). Across all methods, Cheval Corse has the highest Ne/N ratio, while Breton and Comtois have the lowest ratio under the LD and molecular coancestry approach, respectively. This corresponds to a lower breeding population for Cheval Corse (26 individuals) and a much larger breeding population for Breton and Comtois (4368 and 6550, respectively), indicating that a larger census population does not necessarily guarantee a proportionally higher effective population size.

#### Assessment of conservation programs

To evaluate the effectiveness of conservation programs, we analysed how inbreeding and kinship coefficients evolved between the cohorts. We know that, under random mating, kinship coefficients predict the expected inbreeding coefficient for the next generation, while inbreeding coefficients represent the observed value. Additionally, kinship coefficients illustrate the potential for managing genetic diversity, whereas inbreeding coefficients reflect the outcomes of management efforts. Further, to assess whether inbreeding is attributable to recent management or the result of older inbreeding/practices in a closed population, we examined the distribution and mean of the HBD segments.

In Figure 5B, most breeds fall into the first group, characterized by increased inbreeding and kinship coefficients. For all these breeds (except Henson), management and breeding were predominantly regulated by the National Stud Farm before privatization. The conditions in group one suggest a closed breeding program or strong selection after privatization, which is also reflected in the pattern of HBD segments: some breeds in this group showed increased average length and counts above the >16 Mb category (Ardennais, Breton, Percheron, Poitevin Mulassier); some had a decrease in average length but increased counts (Cob Normand, Comtois, Henson, Poney Landais). This could also explain why Poney Landais showed a significant difference in the distribution of HBD segments between the cohorts, as the average across the cohorts remained more or less the same (12.26 Mb versus 14.26 Mb); the change in the proportion of HBD segments altered the overall distribution. Nevertheless, breeds in this category are concerning, and careful management may be required.

Merens and Trait du Nord experienced increased inbreeding while kinship remained unchanged, and their HBD segments showed a decrease in recent average length (>16 Mb) but an increase in count. This suggests that, despite being well managed, as reflected by unchanged kinship, their increase in inbreeding is due to ancestral inbreeding that contributes to the increased HBD counts. In contrast, Camargue showed no change in inbreeding but higher kinship, with a similar average HBD length and counts between the cohorts, while there was a significant difference in the distribution of HBD segments, with a higher range of average HBD segment lengths in cohort II (1.51 to 100.67 Mb) compared to cohort I (1.5 Mb to 73.03 Mb). Similar to Merens and Trait du Nord, Vercors showed an increase in inbreeding but, unlike those breeds with stable kinship, Vercors showed decreased kinship, which is generally the case with matings within isolated groups or with family structures that could be observed in admixture analyses.

Breeds such as Castillonnais (significant decrease in overall average length of HBD segments), Cheval Corse, and Pottok showed lower inbreeding but higher kinship. Looking at the category of HBD higher than >16 Mb, Castillonnais and Cheval Corse showed a sharp decrease across the cohort (34.67 Mb versus 25.2 Mb and 28.7 Mb versus 22.8 Mb, respectively), while Pottok had a similar profile of HBD lengths but a slight decrease in the proportion of HBD counts within the category of 8 Mb to >16 Mb (55% versus 47%). The decrease in recent inbreeding, either through a decrease in average length or count, supports that these breeds have well-managed mating practices and breeding programs.

Boulonnais, on the other hand, despite having a poor genetic diversity profile, as indicated by earlier findings, presents the best-case scenario, with a significantly lower kinship and unchanged inbreeding, which show the potential for improvement with maintained inbreeding. Therefore, looking at both the kinship and inbreeding provides better information on the management efficiency and potential of breed improvement.

## 5. Conclusion

This is the first comprehensive study of French local horse breeds at the genomic scale, investigating their recent history since domestication and genetic management. Using neutral and adaptive population structures of these breeds, we reconstructed their history and, supported by historical records, uncovered several processes they have undergone over time, including selective pressures and outbreeding, to meet specific goals, a pattern common in domestic horses. These results help better understand their evolutionary trajectory. Each breed is distinct in its use, morphology, and geographical origin and can be categorized into two groups: draft and territory. We also examined not only the past of these breeds but also their current status. Thus, we evaluated genetic diversity and assessed management efficiency using a combination of inbreeding and kinship coefficients to monitor changes in genetic diversity due to management changes; while current management practices appear sound, precautions should be taken given their small census and effective population sizes. Finally, our study demonstrates the usefulness and practicality of low-density genomic markers for small local breeds, providing valuable information for their conservation.

## Acknowledgement

We thank the @BRIDGe platform (https://doi.org/10.17180/0791-C793) for DNA extraction and the GENTYANE platform (https://doi.org/10.15454/1.5572409592543596E12) for genotyping. This work was supported by the DivEquid project, funded by the Institut français du cheval et de l’équitation (IFCE; grant no. CS22_10). Yumi Shakya’s doctoral research was funded by the AgroDiv project within the Agroecology and Digital Technologies research program (PEPR Agroécologie et Numérique), supported by French government funding managed by the Agence nationale de la recherche (ANR) under the France 2030 program (grant no. ANR-22-PEAE-0005).

## Supplementary Figures and Tables

## Supplementary Figures and Tables

### Figures

**Figure S1:**
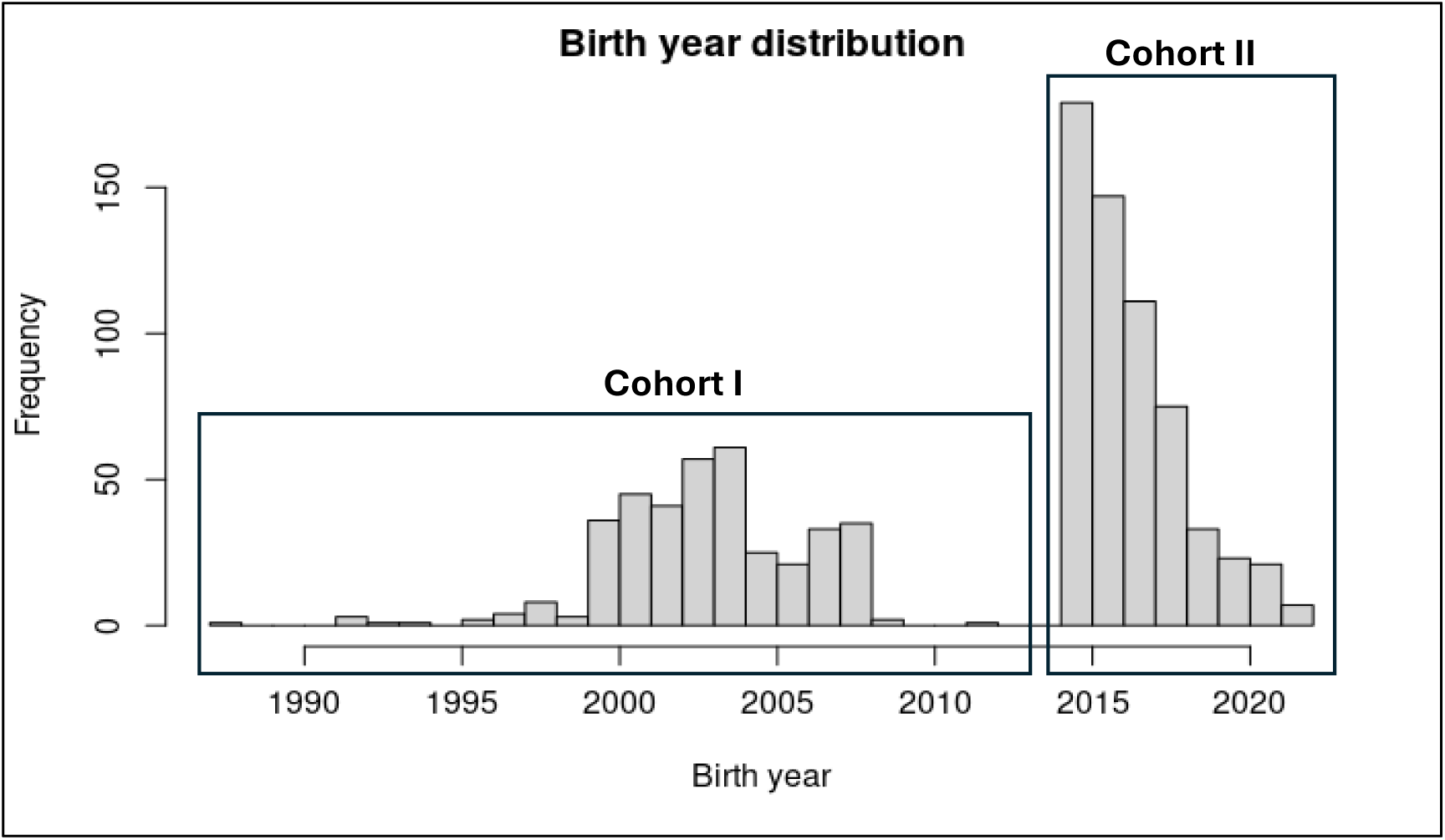
Distribution of samples in two-time cohorts.

**Figure S2:**
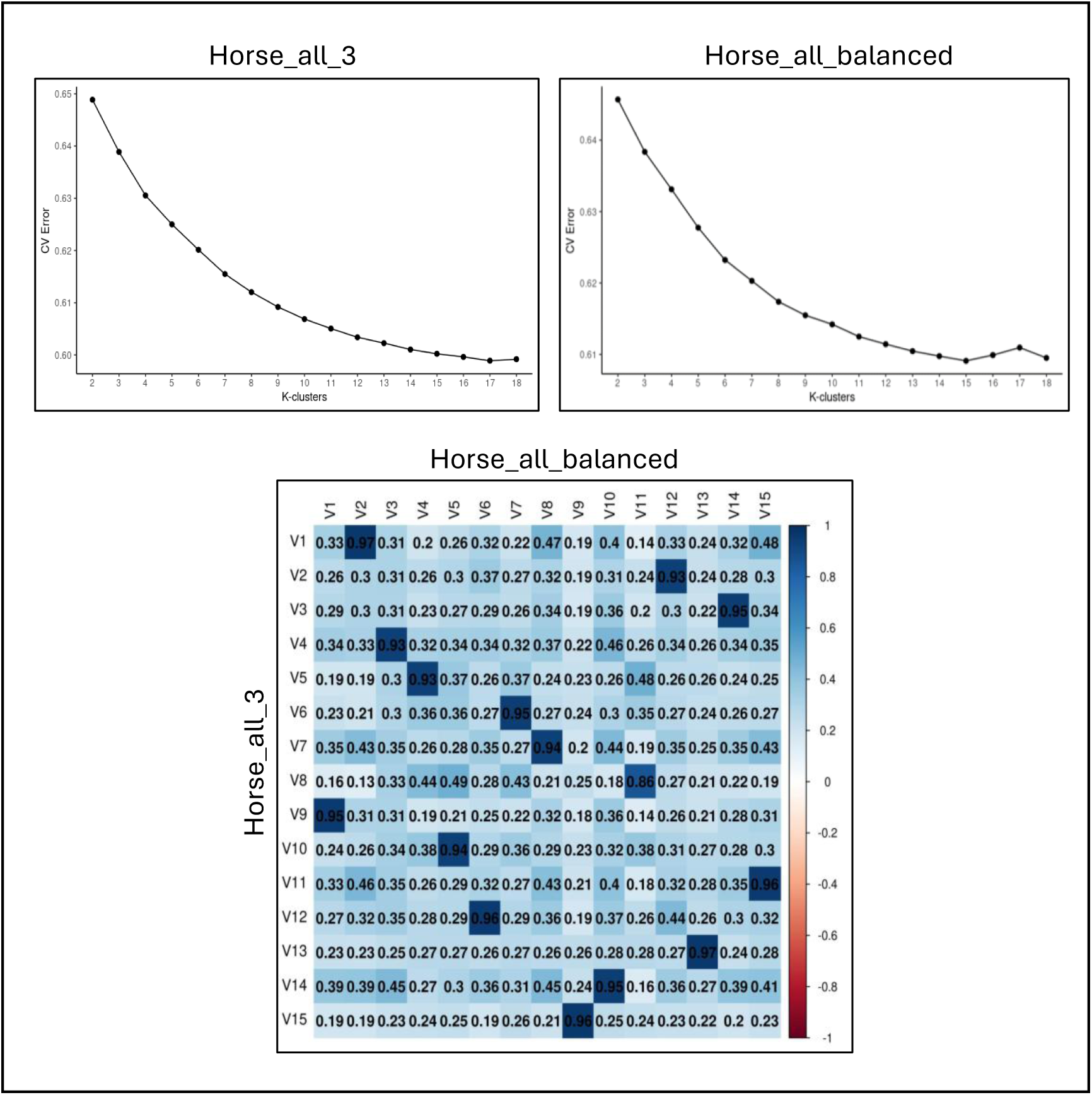
A) Cross-validation errors obtained from the complete dataset, B) balanced dataset, with k from 2 to 18. C) Correlation between ancestry allele frequency matrices at K=15 for both balanced and complete datasets.

**Figure S3:**
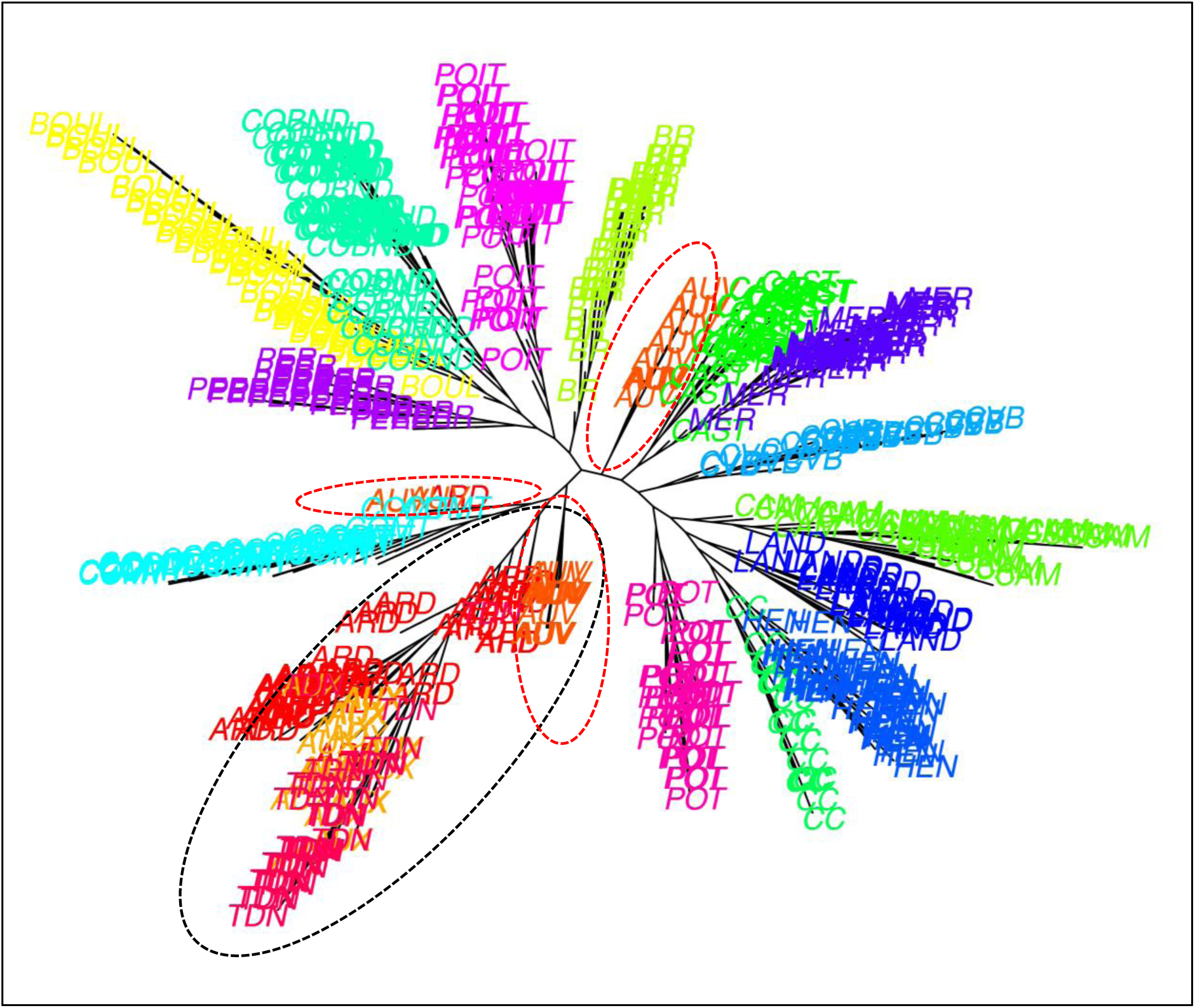
Unrooted neighbour-joining tree obtained from identity by state (IBS) matrix for individuals of 18 breeds, each colour representing a breed, red dotted circle represents three AUV clusters, and black dotted circle represents the joint cluster of ARD, TDN, and AUX.

**Figure S4:**
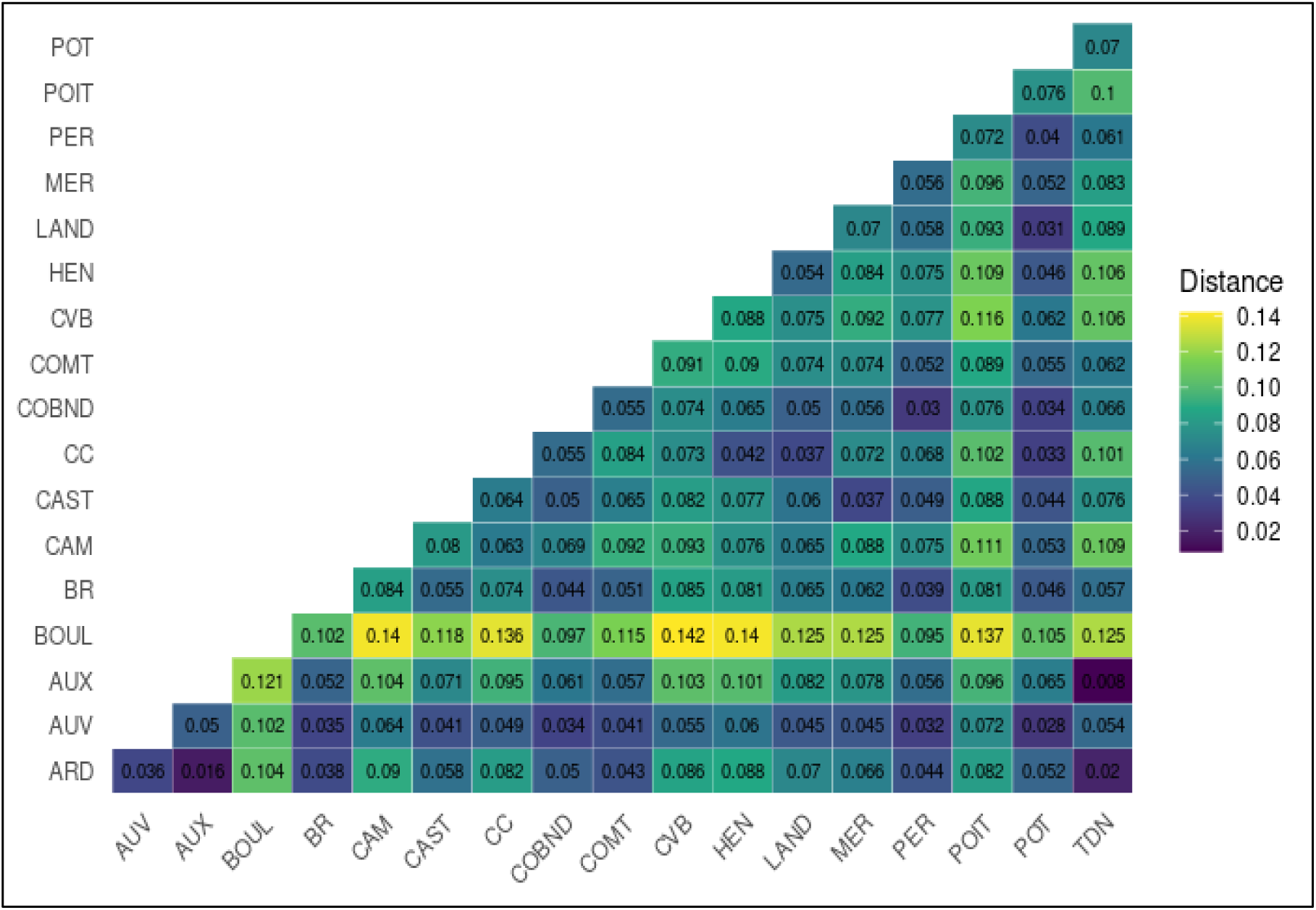
Heatmap representing pairwise F_ST v_alues for 18 breeds.

**Figure S5:**
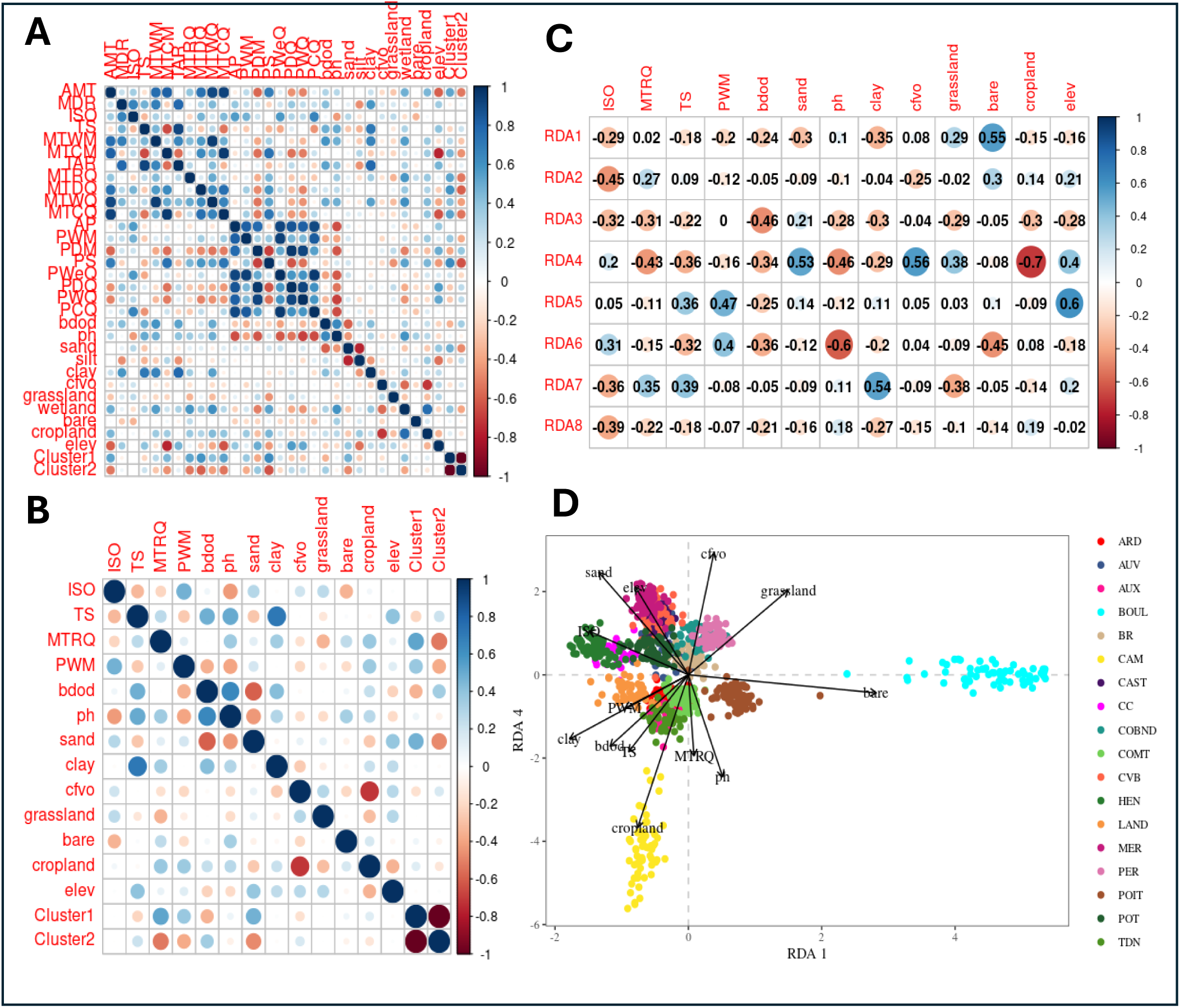
A) Correlation between all the climatic, elevation, soil, landcover and cluster variables. B) Correlation between the environmental variables (4-climatic, elevation, 5-soil, 3-land coverage and 2-clusters) selected after backward variable selection. C) Correlation of selected variables with the first eight RDA axes. D) RDA biplot, the results of the genotype-environment association using partial RDA (environmental model) for RDA1 versus RDA4. Variable abbreviations are given in Table 4.

**Figure S6:**
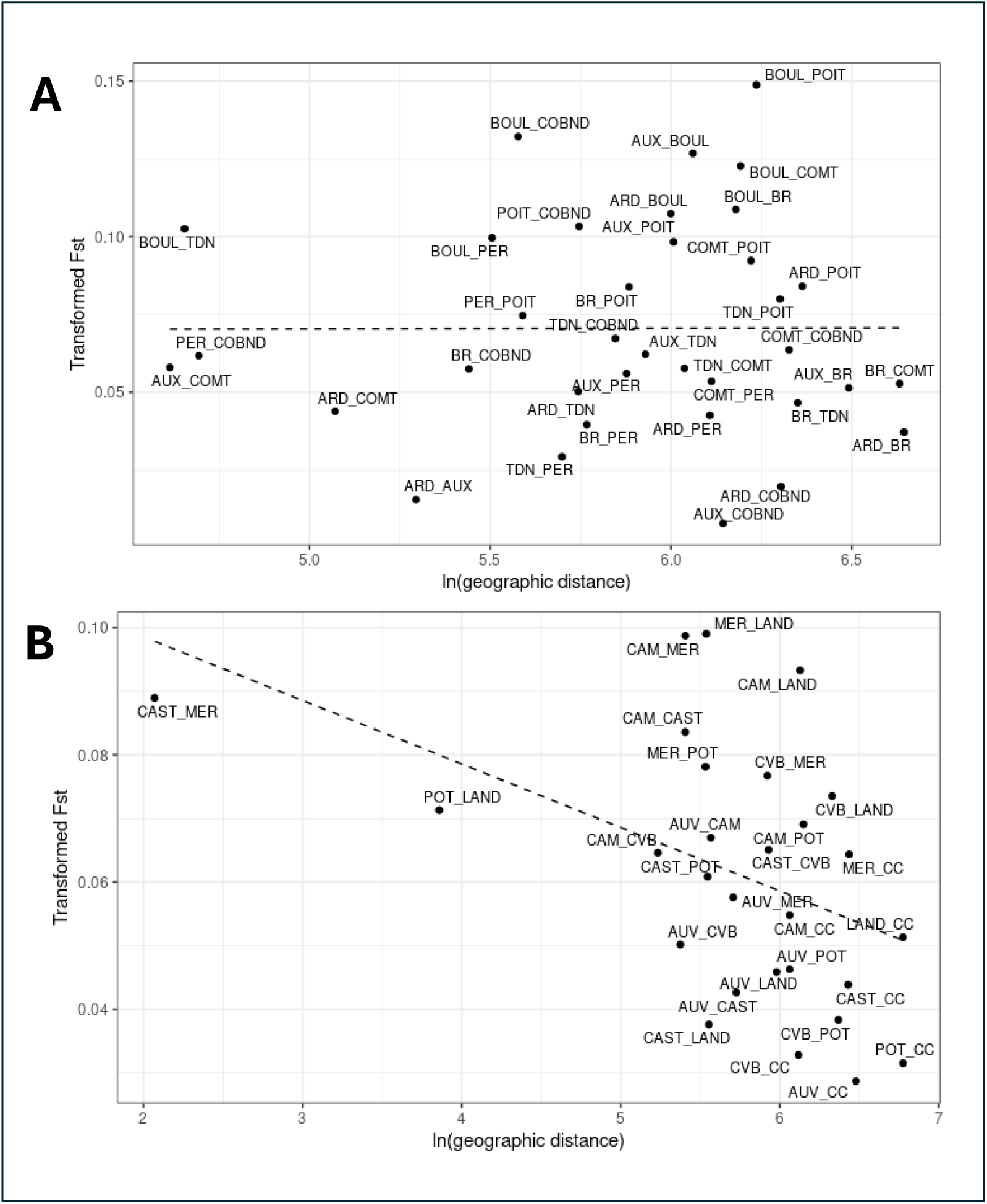
A) Mantel test for nine draft breeds. B) Mantel test for eight territory breeds. Breed abbreviations are given in Table 1.

**Figure S7:**
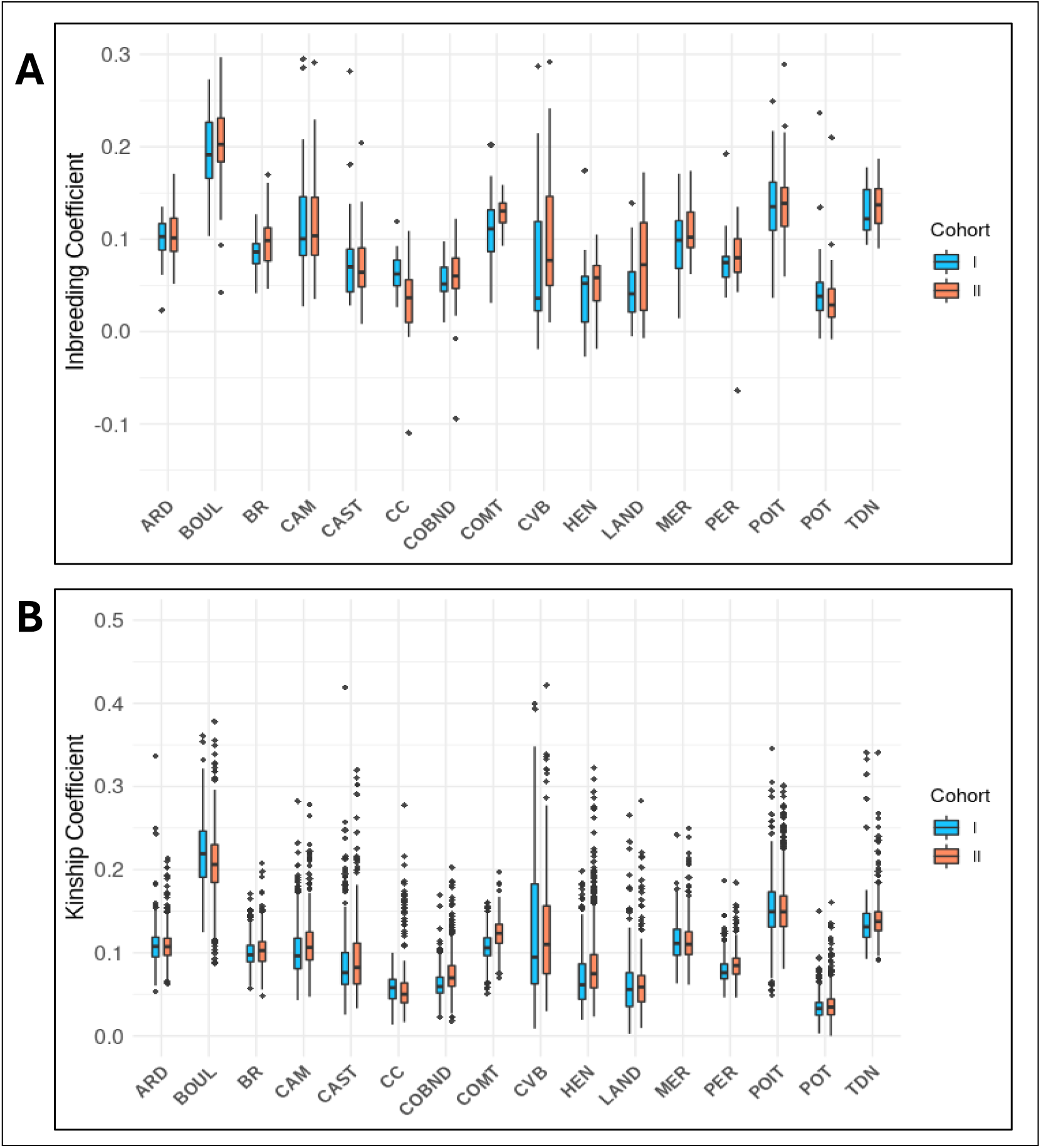
Distribution of A) Inbreeding coefficients and B) Kinship coefficients obtained from the POPKIN R package in 16 breeds (excluding AUV and AUX) in two time cohorts.

### Tables

**Table S1:**
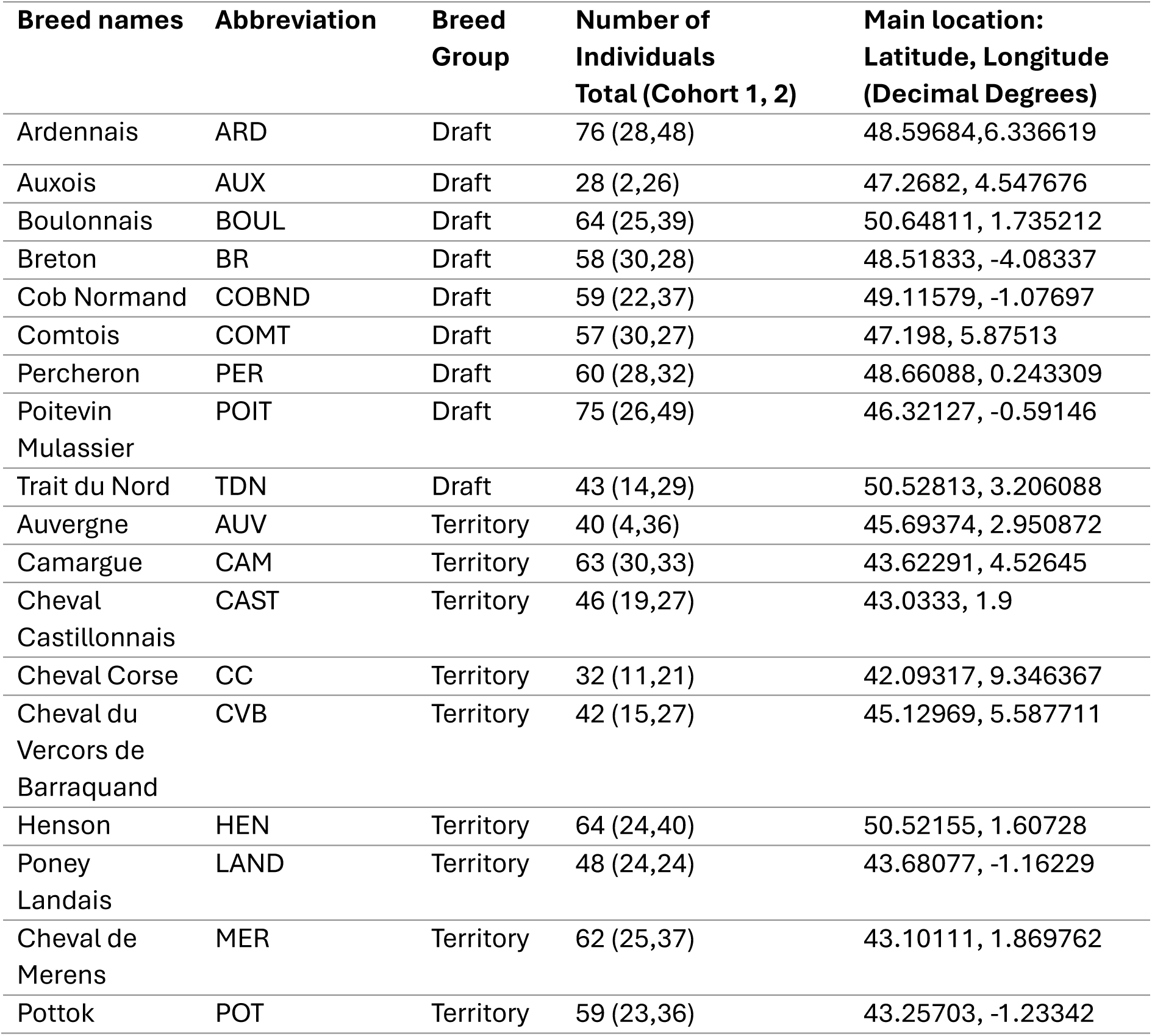
Information on horse breeds used in the study.

**Table S2:**
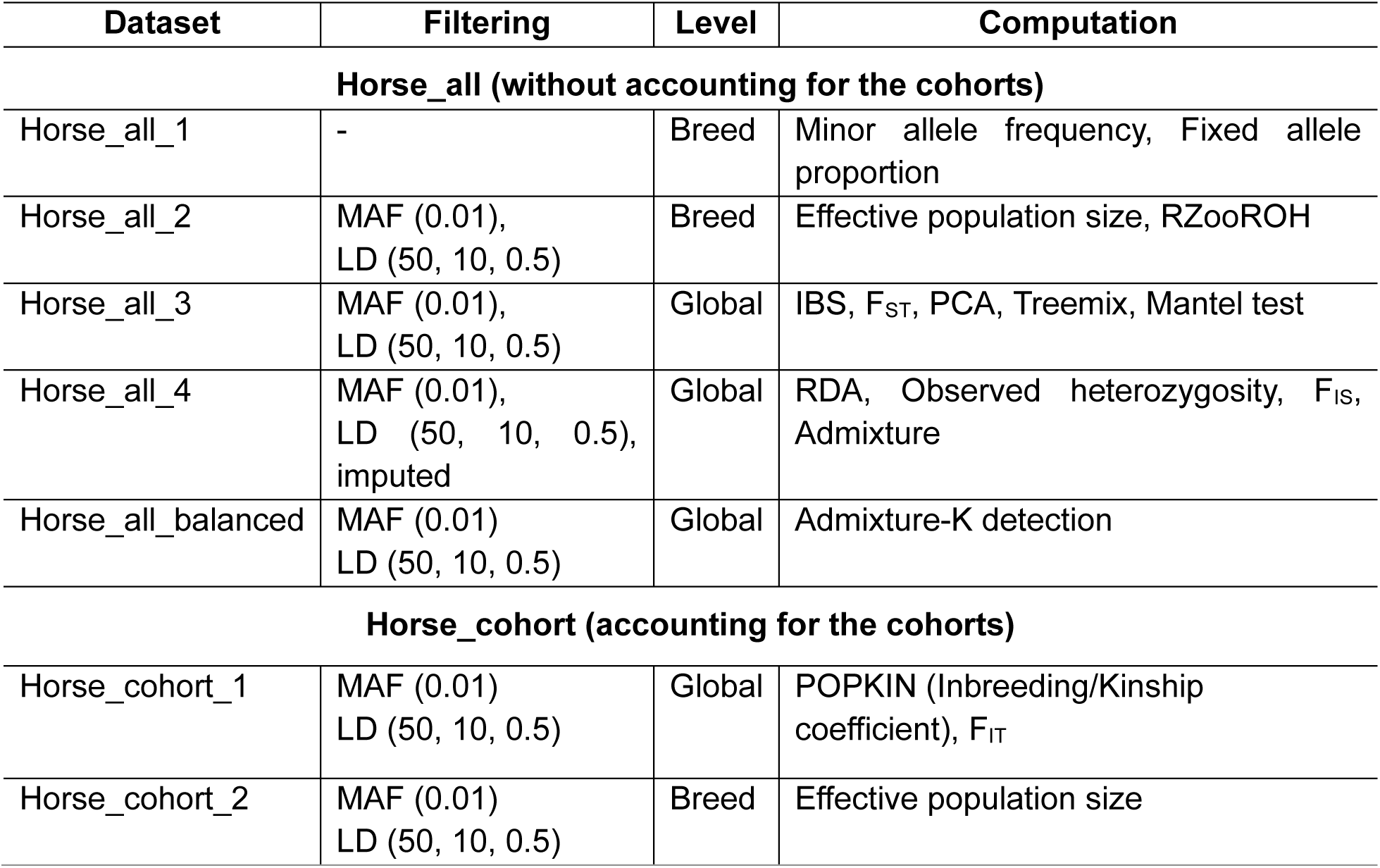
List of datasets, filter parameters, and type of analyses.

**Table S3:**
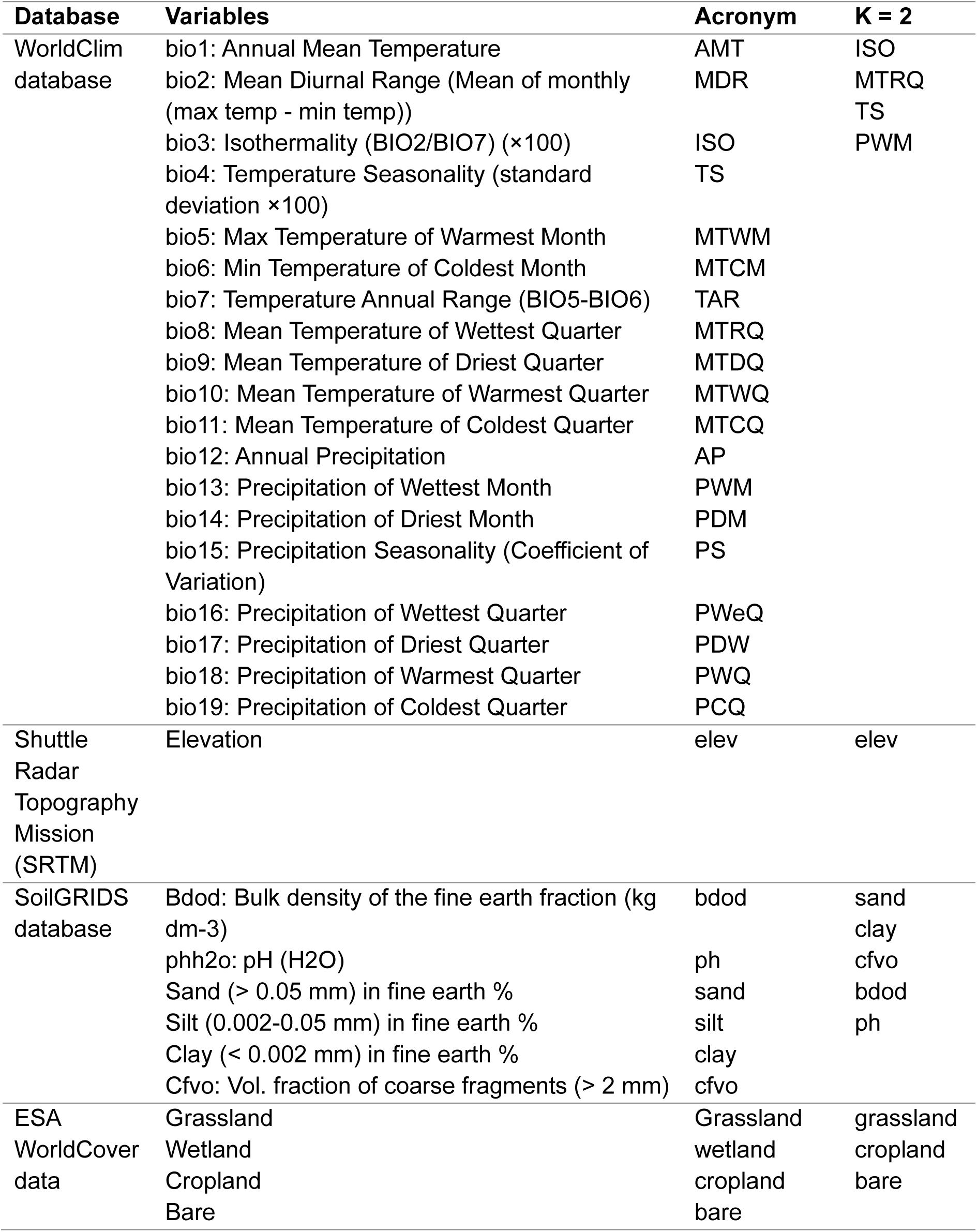
List of climatic, elevation, soil, and landcover variables before and after backward variable selection.

**Table S4:**
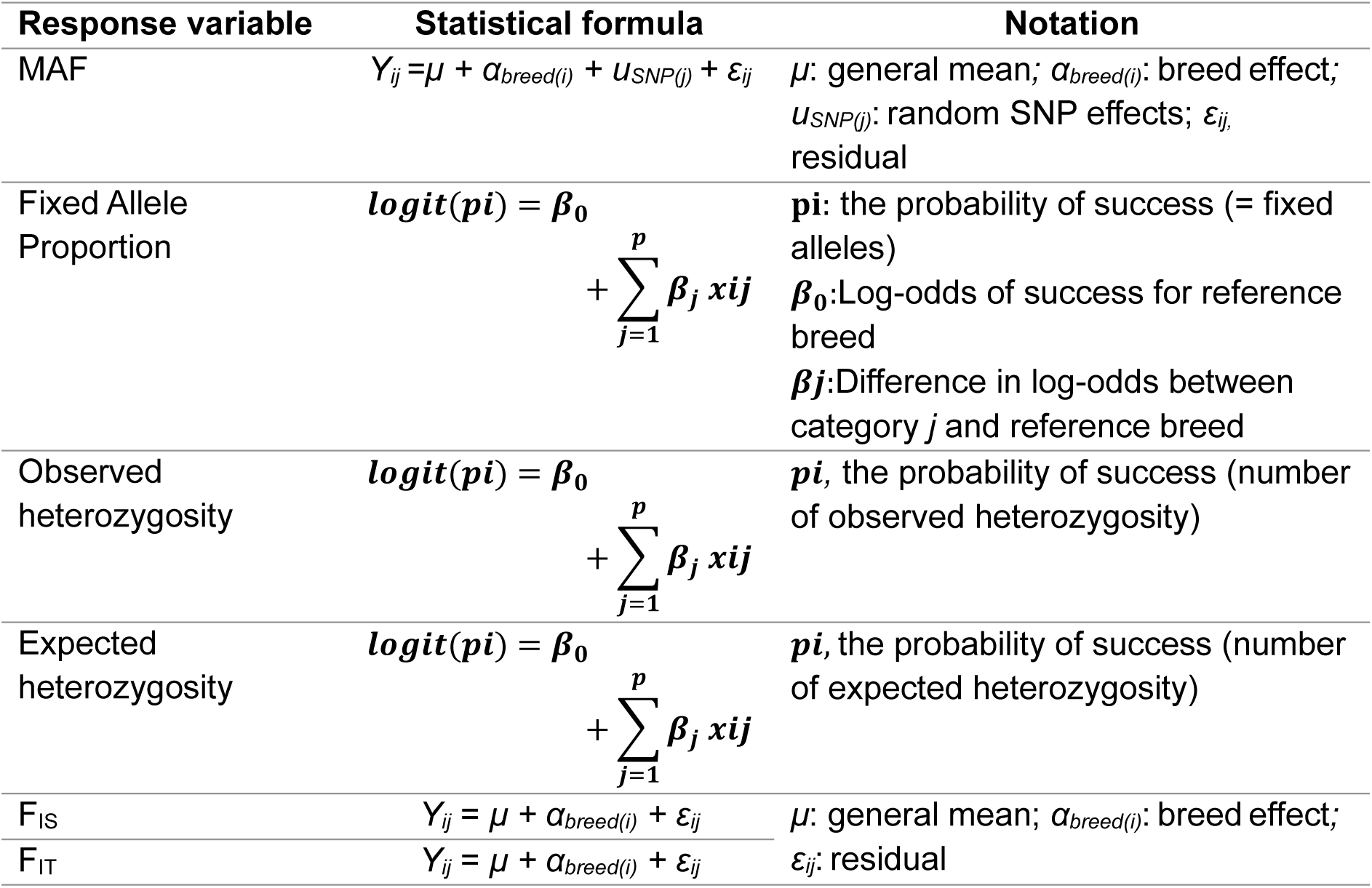
Statistical formulas for comparing diversity parameters among different breeds.

**Table S5:**
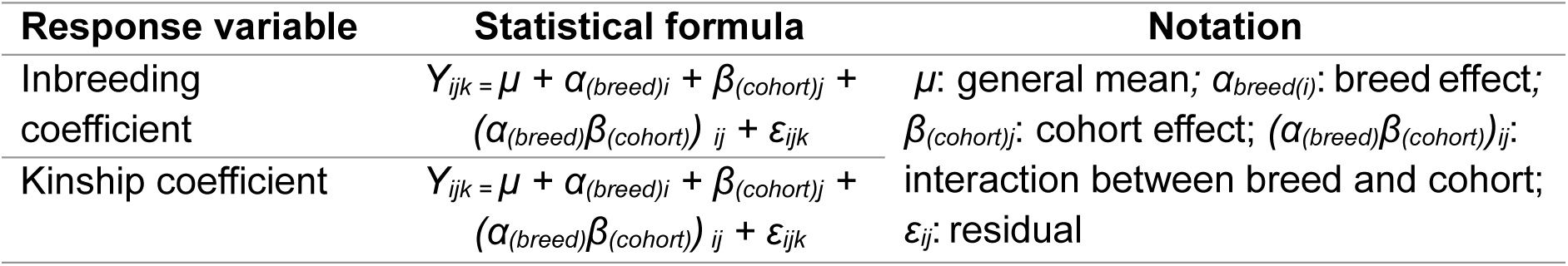
Statistical formula for comparing inbreeding and kinship coefficients across breeds and cohorts.

**Table S6:** Effective Population Size (Ne), Total Breeding Population (N), and the Ratio of Ne/N.

| Breeds | LD | Het | Mol | Sex-ratio | Breeding population(N) | LD/N | het/N | Mol/N | sex-ratio/N |
| --- | --- | --- | --- | --- | --- | --- | --- | --- | --- |
| ARD | 303.6 | 54.3 | 5.5 | 525.338 | 1118 | 0.27 | 0.05 | 0.005 | 0.47 |
| AUV | 44 | 27.2 | 15.2 | 57.679 | 112 | 0.39 | 0.24 | 0.136 | 0.52 |
| AUX | 99.8 | 27 | 16.7 | 87.562 | 201 | 0.49 | 0.13 | 0.083 | 0.44 |
| BOUL | 129.8 | 24.9 | 1.8 | 129.987 | 304 | 0.43 | 0.08 | 0.006 | 0.43 |
| BR | 297.1 | 31.4 | 14.7 | 1823.23 | 4368 | 0.07 | 0.007 | 0.003 | 0.42 |
| CAM | 127.2 | NA | 6.4 | 639.158 | 1026 | 0.12 | NA | 0.006 | 0.62 |
| CAST | 36.8 | 19.3 | 7.1 | 36.706 | 51 | 0.72 | 0.38 | 0.139 | 0.72 |
| CC | 58.4 | 15.9 | 11 | 20.462 | 26 | 2.25 | 0.61 | 0.423 | 0.79 |
| COBND | 201.2 | 34 | 12.1 | 139.862 | 377 | 0.53 | 0.09 | 0.032 | 0.37 |
| COMT | 524.9 | NA | 4 | 2481.871 | 6550 | 0.08 | NA | 0.001 | 0.38 |
| CVB | 22.8 | 11.8 | 2.1 | 23.1 | 40 | 0.57 | 0.29 | 0.053 | 0.58 |
| HEN | 59 | 16.2 | 11.7 | 44.047 | 85 | 0.69 | 0.19 | 0.138 | 0.52 |
| LAND | 61.7 | 40 | 7 | 76.408 | 98 | 0.63 | 0.41 | 0.071 | 0.78 |
| MER | 221 | 39.3 | 10.4 | 184.938 | 469 | 0.47 | 0.08 | 0.022 | 0.39 |
| PER | 284.5 | 79.5 | 19.5 | 563.069 | 1688 | 0.17 | 0.05 | 0.012 | 0.33 |
| POIT | 112.5 | 26.7 | 2.4 | 106.376 | 170 | 0.66 | 0.16 | 0.014 | 0.63 |
| POT | 262.4 | NA | 19.3 | 345.782 | 1055 | 0.25 | NA | 0.018 | 0.33 |
| TDN | 154.4 | 33.6 | 11.1 | 29.919 | 123 | 1.26 | 0.27 | 0.09 | 0.24 |
| **Mean** | 166.728 | 32.073 | 9.889 | 406.416 | 992.278 | 0.56 | 0.20 | 0.07 | 0.49 |

*LD: Linkage disequilibrium, Het: Heterozygous excess, Mol: Molecular Coancestry, Sex-ratio: equation from Wright (1938) (Formula given in the main text)*

**Table S7:**
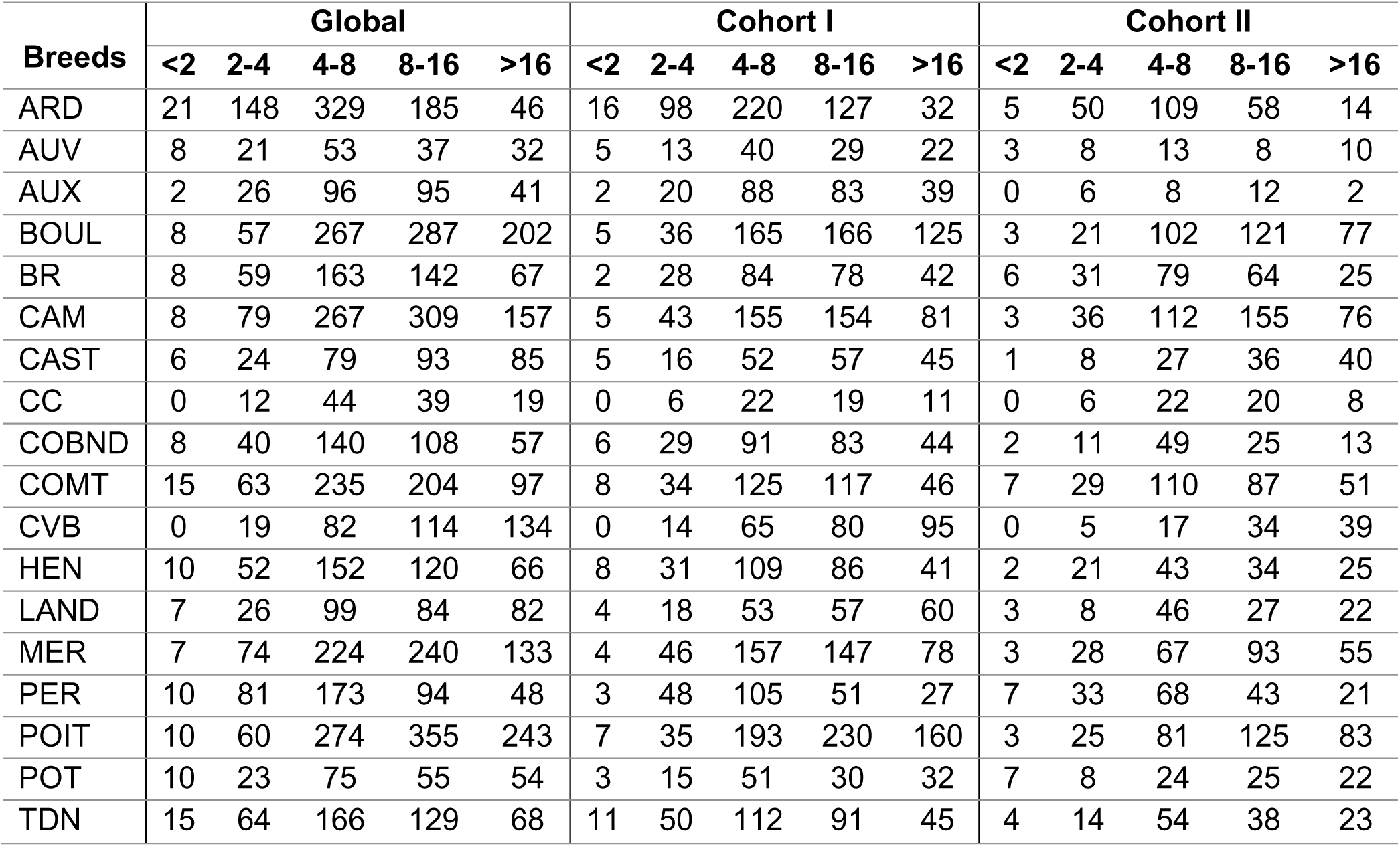
Number of HBD segments detected per class (Mb) at a global and cohort level.

**Table S8:** Average length of HBD segments with minimum and maximum values at a global and cohort level.

| **Breeds** | **Global** | **min** | **max** | **Cohort I** | **min** | **max** | **Cohort II** | **min** | **max** |
| --- | --- | --- | --- | --- | --- | --- | --- | --- | --- |
| ARD | 7.54 | 0.13 | 59 | 7.29 | 1.31 | 51 | 7.65 | 0.134 | 58.96 |
| AUV | 11.6 | 0.25 | 61.9 | 11.27 | 0.25 | 49 | 11.72 | 0.51 | 61.88 |
| AUX | 10.33 | 1.58 | 45.3 | 9.35 | 2.55 | 25 | 10.44 | 1.575 | 45.31 |
| BOUL | 12.06 | 0.54 | 83.8 | 11.94 | 0.81 | 54 | 12.13 | 0.539 | 83.79 |
| BR | 9.708 | 0.95 | 54.4 | 8.97 | 0.95 | 30 | 10.35 | 1.453 | 54.37 |
| CAM | 11.84 | 1.51 | 101 | 12.21 | 1.54 | 73 | 11.51 | 1.511 | 100.7 |
| CAST | 14.61 | 0.81 | 88.2 | 17.75 | 1.86 | 88 | 12.6 | 0.814 | 49.75 |
| CC | 10.48 | 2.16 | 76.2 | 10.66 | 2.16 | 76 | 10.29 | 2.612 | 33.71 |
| COBND | 10.57 | 0.3 | 75.6 | 10.5 | 0.3 | 76 | 10.58 | 0.555 | 60.4 |
| COMT | 10.16 | 0.48 | 67.6 | 10.65 | 0.94 | 68 | 9.72 | 0.481 | 49.21 |
| CVB | 16.3 | 2.05 | 98.2 | 16.77 | 2.05 | 59 | 16.11 | 2.051 | 98.21 |
| HEN | 9.92 | 0.85 | 63.2 | 10.93 | 1.54 | 58 | 9.45 | 0.845 | 63.21 |
| LAND | 13.56 | 1.16 | 75 | 12.26 | 1.16 | 62 | 14.26 | 1.251 | 75.05 |
| MER | 11.48 | 1.04 | 88.9 | 12.41 | 1.1 | 57 | 10.95 | 1.041 | 88.88 |
| PER | 8.729 | 0.31 | 78.2 | 8.65 | 0.31 | 38 | 8.78 | 1.02 | 78.18 |
| POIT | 12.77 | 1.54 | 82.7 | 12.55 | 1.54 | 64 | 12.87 | 1.589 | 82.74 |
| POT | 13.02 | 0.4 | 84.4 | 13.21 | 0.4 | 84 | 12.9 | 0.553 | 63.89 |
| TDN | 9.714 | 1.1 | 58.8 | 10.58 | 1.53 | 48 | 9.34 | 1.102 | 58.75 |

